# Differing components of plasticity in threshold traits cause diverging eco-evolutionary responses to spatio-seasonal environmental deterioration

**DOI:** 10.64898/2026.09.09.750077

**Authors:** Thomas R. Haaland, Paul Acker, Ana Payo-Payo, Jane M. Reid

**Affiliations:** Department of Biology, Norwegian University of Science and Technology, Trondheim, Norway; Departamente Biología Animal, Parasitología, Ecología, Edafología y Química Agrícola, Universidad de Salamanca, Salamanca, Spain; School of Biological Sciences, University of Aberdeen, Aberdeen, United Kingdom

**Keywords:** developmental plasticity, cohort effect, threshold trait, seasonal migration, evolutionary rescue, frequency-dependence

## Abstract

Eco-evolutionary responses to long-term environmental deteriorations will fundamentally depend on interactions between plasticity and evolution of life-history traits that shape population dynamics. Counter-intuitive evolutionary and (meta)population dynamics could arise when different components of individual-specific and/or shared site-specific developmental and labile plasticity affect traits with intrinsically non-linear genotype-environment-phenotype relationships, especially given density-dependent fitness outcomes. Frequency-dependent evolutionary responses could then emerge, but resulting eco-evolutionary dynamics and outcomes have rarely been considered. By modelling a partially-migratory metapopulation encompassing facultative seasonal migration versus residence formulated as a quantitative genetic threshold trait, we show how different forms of plasticity in liability to migrate interact with spatio-seasonal metapopulation dynamics to generate divergent eco-evolutionary responses to spatially restricted environmental deterioration. Temporary and permanent individual-specific environmental effects induced faster evolutionary recovery than might simply be expected, by revealing cryptic genetic variation and allowing adaptive phenotypic changes through repeated episodes of selective disappearance. Conversely, shared subpopulation-specific environmental effects caused among-year variation in phenotype frequencies, impeding evolutionary responses to the extent that migratory metapopulation connectivity was ultimately eradicated. We thereby reveal key principles of how structurally different forms of plasticity in a dichotomous quantitative genetic trait can induce complex eco-evolutionary dynamics, culminating in differing degrees of evolutionary rescue versus constraint.

## Introduction

Major contemporary ambitions in evolutionary ecology are to understand joint evolutionary, phenotypic and population dynamic responses to deteriorating environmental conditions, and thereby predict potentials for evolutionary rescue versus population collapse (Brunner et al. 2019, Fouqueau and Polechová 2024, Urban et al. 2024). These ambitions require integrating genetic and environmental effects on key phenotypes and resulting demography, and hence predicting the progress of rapid adaptive evolution alongside manifestations of phenotypic plasticity in the context of changing environments and population sizes (Chevin et al. 2010, Merilä and Hendry 2014, Urban et al. 2024). Here, considerable empirical and theoretical work has highlighted the potentially interacting roles of evolution and plasticity in overcoming rapid environmental changes (Parmesan 2006, Bell and Collins 2008, Reed et al. 2010, Carlson et al. 2014, Ashander et al. 2016, Lane et al. 2018, Crowther et al. 2023, Martin et al. 2023). Not least, adaptive plasticity could delay extinction, yet slow evolutionary responses by decreasing heritability and/or weakening selection, thereby delaying adaptive evolution (Price et al. 2003, Paenke et al. 2007, Chevin et al. 2010, Oostra et al. 2018, Muñoz 2021, Fouqueau and Polechová 2024). However, eco-evolutionary theory has not yet considered realistically complex situations where key phenotypes are shaped by multiple spatially and temporally varying stochastic environmental effects and resulting forms of plasticity, given intrinsically non-linear genotype-environment-phenotype relationships and multi-year life-histories (Snell-Rood et al. 2018, Govaert et al. 2019, Klausmeier et al. 2020, Yamamichi 2022, Urban et al. 2024). Such effects could generate complex eco-evolutionary feedbacks with non-intuitive outcomes, particularly when affecting life-history traits that also experience density-dependent and/or frequency-dependent selection (De Meester et al. 2019, Lion et al. 2022).

Accounting for multi-faceted plastic effects is necessary because empirical studies of life-history traits often identify multiple substantial components of phenotypic variation that are attributed to known and/or unknown current or previous environmental conditions (Kendall et al. 2011, Charmantier et al. 2014, Cam et al. 2016, Forsythe et al. 2021). These components can be shared by sets of individuals that experience common environments in space and/or time (generating site-, year-or cohort-specific effects), or be unique to individuals (individual-specific effects). Such effects can be effectively life-long permanent, stemming from ‘developmental’ plasticity in response to early-life conditions, or temporary, representing ‘labile’ plasticity in response to current conditions. These structurally differing forms of plasticity have very different implications. For example, permanent individual-specific effects could show protracted responses to selection, where selective disappearance causes lasting changes in population mean phenotype even without any underlying genetic variation or hence evolution (Cam et al. 2002, van de Pol and Verhulst 2006, Wynn et al. 2025). In contrast, shared site-, year-or cohort-specific effects could align the phenotypes of sets of individuals, generating less concurrent among-individual variation and hence less opportunity for selection. Meanwhile, selection on temporary effects that are continuously regenerated will have no direct lasting phenotypic effect, while such effects could partially mask additive genetic variation and hence slow evolution (effectively reducing heritability). Yet, despite these potential impacts, the degree to which differing forms of environmental effects can reshape evolutionary, population dynamic and eco-evolutionary responses to environmental changes has scarcely been considered (Lindström and Kokko 2002, Vindenes et al. 2008, Lyberger et al. 2021, Forsythe et al. 2024).

Resulting dynamics will be further complicated given traits that are expressed as discrete alternative phenotypes, implying highly non-linear genotype-environment-phenotype relationships. Such traits are commonplace, encompassing behavior, morphology and life-history, and are often appropriately conceptualised as quantitative genetic ‘threshold traits’ where alternative dichotomous phenotypes are expressed when an underlying continuous, unbounded ‘liability’ is above versus below some threshold (Roff 1996, Pulido 2011, Dodson et al. 2013, Berg et al. 2019, Reid and Acker 2022, Acker and Reid 2025). This liability can comprise combinations of genetic and multiple environmental components, including temporary or permanent individual-, site-, year-and/or cohort-specific effects. Multiple intrinsic properties of such threshold traits could then substantially affect phenotypic and evolutionary dynamics. Notably, evolutionary responses to selection are frequency-dependent, decreasing as phenotype frequencies deviate further from 0.5 (Dempster and Lerner 1950, Roff 1996). Consequently, shared site-, year-and/or cohort-specific environmental effects on liability could directly alter rates of evolution because they affect phenotype frequencies. Further, threshold traits can harbor considerable ‘cryptic’ genetic variation in liability, which is only phenotypically expressed given large environmental deviations (Suzuki and Nijhout 2006, McGuigan and Sgrò 2009). Intrinsic gene-by-environment interactions then emerge on the phenotypic scale, even if genetic and environmental effects on the underlying liability are strictly additive, depending on whether the mean liability is near or far from the threshold (Reid and Acker 2022). The degree to which cryptic genetic variation is temporarily exposed to selection, potentially inducing evolution, will thus depend on the form and magnitude of environmental effects (i.e. of plasticity).

Yet, despite their broad relevance, these intrinsic properties of dichotomous threshold traits, and their connections with plasticity, have scarcely been considered in efforts to evaluate eco-evolutionary responses to environmental deterioration. Most models instead envisage traits that are continuously distributed on observed phenotypic scales with linear genotype-environment-phenotype relationships, implying unbounded evolution that can be readily predicted as products of selection gradients and additive genetic variances (Bürger and Lynch 1995, Chevin et al. 2010, Ashander et al. 2016, Scheiner et al. 2020, but see Khare et al. 2024). Furthermore, most models envisage short-lived organisms with simple life-cycles and non-overlapping cohorts and generations, largely eliminating distinctions between developmental and labile plasticity (Gomulkiewicz and Holt 1995, Lande 2009, Lindsey et al. 2013, Carlson et al. 2014, Orr and Unckless 2014, Bell 2017). Such models consequently cannot capture eco-evolutionary dynamics involving interactions among evolution, selection and multiple forms of plasticity that could commonly arise in longer-lived organisms. New eco-evolutionary models that explicitly incorporate threshold trait dynamics and multi-year life-histories could therefore reveal new properties of combined plastic and evolutionary responses to environmental deterioration (de Zoeten and Pulido 2020), including the roles of different types of environmental effects in driving evolutionary rescue or extinction (e.g. Gomulkiewicz and Shaw 2013, Klausmeier et al. 2020, Crowther et al. 2023).

One ecologically critical setting, which epitomizes the limitations of existing eco-evolutionary models, concerns facultative dichotomous expression of reversible seasonal migration versus year-round residence in seasonally-varying environments (Milner-Gulland et al. 2011, Bauer and Hoye 2014, Reid et al. 2018). Here, liability to migrate, and resulting phenotypic expression of seasonal migration versus residence (representing a threshold trait), is commonly heritable but also shows substantial permanent environmental variation (i.e. developmental plasticity), generating high adult phenotypic repeatabilities and strong cohort effects, alongside labile plasticity (observed in partially-migratory birds, fish and mammals, Pulido 2007, Dodson et al. 2013, Eggeman et al. 2016, Chambon et al. 2019, Acker et al. 2023, Ugland et al. 2026). Importantly, the expressed degree of seasonal migration directly shapes spatio-seasonal population dynamics (i.e. variation in numbers and seasonal distributions of surviving individuals, Payo-Payo et al. 2022), in turn shaping the form and strength of selection caused by any local environmental deterioration. Thus, environmental and genetic effects on seasonal migration will fundamentally determine how combinations of plasticity and evolution drive spatio-seasonal eco-evolutionary dynamics (Reid et al. 2018, Haaland et al. 2026).

Further interlinked dynamics could then arise within ‘partially migratory metapopulations’ (PMMPs), where spatially discrete subpopulations (i.e., groups of individuals breeding together) overlap with other subpopulations during non-breeding seasons, depending on their degrees and forms of seasonal migration versus residence (e.g. Figure 1A; Papastamatiou et al. 2013, Geijer et al. 2016, Reid et al. 2018, Allen et al. 2019). Deteriorating non-breeding season conditions in one location could then induce evolutionary changes in seasonal migration that directly and indirectly affect multiple subpopulations throughout the PMMP (Reid et al. 2018, Haaland et al. 2026). For example, evolutionary rescue in one subpopulation, manifested as subpopulation decline due to environmental deterioration followed by evolutionary recovery, could reduce other subpopulation sizes. This could occur because evolutionary and/or plastic increases in seasonal migration increase non-breeding season densities elsewhere, hence reducing density-dependent survival (Taylor and Norris 2010, Haaland et al. 2026). Such density-dependence can thereby generate frequency-dependent selection, which could contribute to maintaining phenotypic variation (Lundberg 1988, Kaitala et al. 1993). However, eco-evolutionary dynamics resulting from interacting combinations of multi-faceted plasticity, non-linear genotype-environment-phenotype relationships and density-dependence, have not been examined.

**Figure 1:**
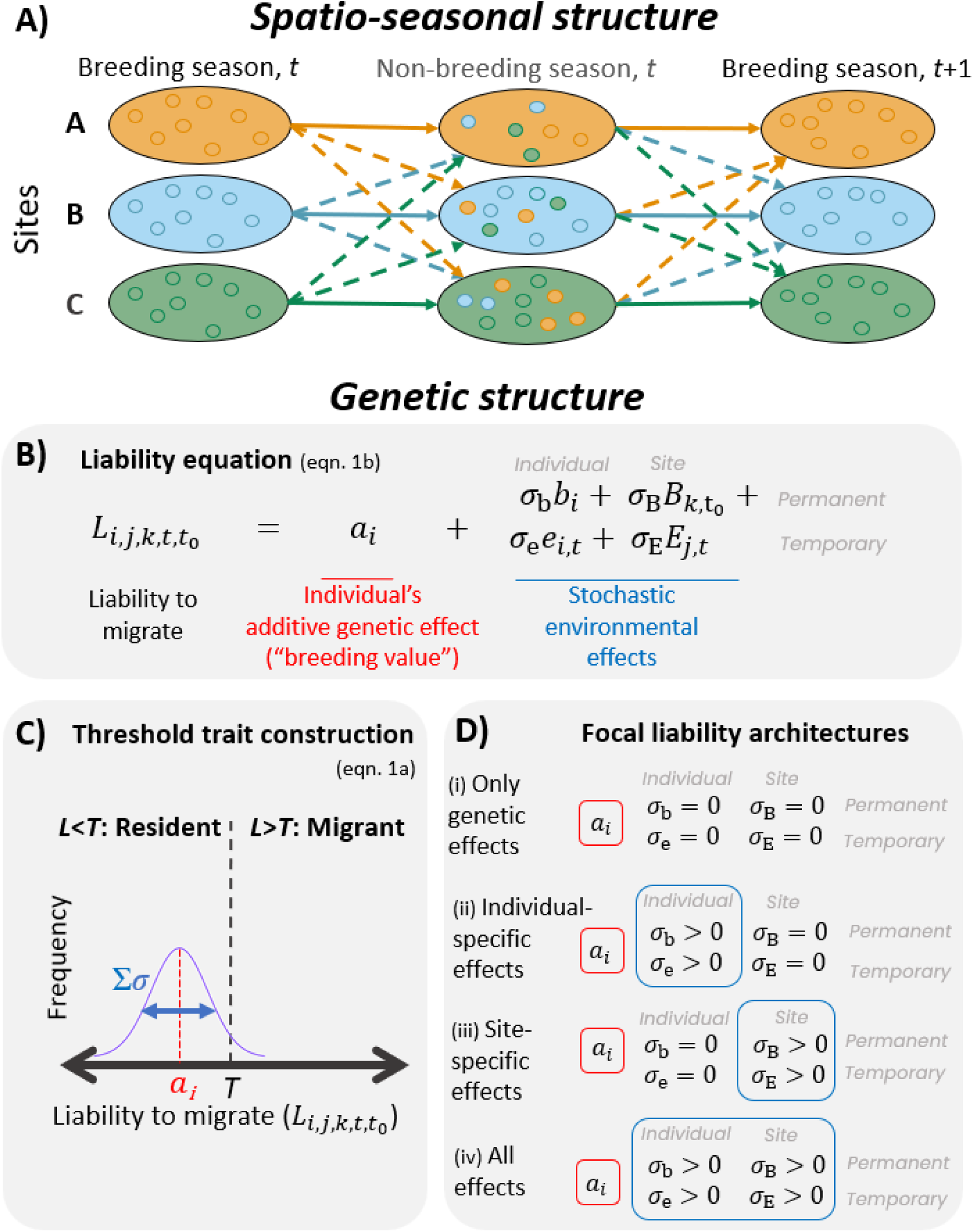
Overview of the modelled spatio-seasonal and genetic structures. (A) Illustration of *N*=3 sites (large colored ovals) harboring partially-migratory subpopulations of individuals (filled circles of corresponding colors) that can remain resident in their home site year-round, or temporarily migrate to a different site for the non-breeding season. Solid and dashed arrows respectively denote residence and migration, whose frequencies could vary substantially within and among sites. (B) Liability to migrate, *L_i_*_,*j*,*k*,*t*,*t*_0 (eqn. 1b), can consist of additive genetic (red) and multiple stochastic environmental (blue) effects, where *b*, *B*, *e*, *E* are standardized environmental effects (z-scores), and *σ*-parameters represent the magnitudes of deviations. (C) Illustration of the quantitative genetic threshold trait structure, where individuals migrate if their liability *L_i_*_,*j*,*k*,*t*,*t*_0 exceeds the threshold *T* (eqn. 1a). An individual’s additive genetic effect *ai* effectively represents its liability in mean environmental conditions. Stochastic environmental effects cause liability distributions with widths determined by the *σ*-parameters. (D) Environmental effects can take any combination of permanent or temporary and individual-specific or site-specific, with four focal liability architectures (i-iv). Subscripts refer to: *i* – individual; *j* – breeding site; *k* – birth site; *t* – current year; *t*_0_ – birth year.

Accordingly, we tested how different forms of environmental effects (and hence of plasticity) can shape eco-evolutionary dynamics in spatially and seasonally varying environments given intrinsically non-linear genotype-environment-phenotype relationships underlying the ecologically critical trait of facultative seasonal migration. To achieve this, we modelled a PMMP with seasonal migration versus residence formulated as a quantitative genetic threshold trait, with the underlying liability determined by combinations of additive genetic and permanent and temporary environmental effects given a multi-year life-history. By imposing gradual deterioration in local non-breeding season environmental conditions we test the degree to which rapid evolution of seasonal migration can ‘rescue’ subpopulations; whether such rescue is facilitated or impeded by different forms of environmental effects on liability; and how local evolution of seasonal migration percolates throughout the PMMP through spatially and seasonally structured density-dependence to affect evolution of migration and resulting population dynamics elsewhere. We reveal how these outcomes are generated by interacting dynamics of genetic, environmental and phenotypic effects and resulting selection, when evolution is allowed versus prohibited. We thereby show how counter-intuitive and non-linear evolutionary and phenotypic dynamics induced by underlying plasticity can arise and cause simultaneous and interacting positive and negative effects on the dynamics of spatio-seasonally interlinked subpopulations.

## Methods

To capture spatio-seasonal eco-evolutionary dynamics emerging within PMMPs given systematic local environmental deterioration alongside multiple components of stochastic environmental variation and density-dependence, we built a spatially and seasonally structured quantitative genetic individual-based model (Reid et al. 2018, Govaert et al. 2019). Our model tracks individuals’ survival, reproduction and movements across sequential breeding and non-breeding seasons, linked by episodes of potential seasonal migration (Haaland et al. 2026). Individuals breeding within each of *N* sites, which can differ in seasonal habitat quality, can either remain resident at their breeding site year-round, or migrate (i.e. reversibly move) to a different site for the non-breeding season before returning to their breeding site for the following breeding season (Fig. 1A). Each site therefore harbors distinct subpopulations that are spatially separated during the breeding season, but may partially overlap during the non-breeding season. Our model can be parameterized to envisage diverse partially migratory systems, including where seasonal migrations span altitudinal, latitudinal and/or hydrological gradients (as observed in diverse taxa, Papastamatiou et al. 2013, Geijer et al. 2016, Allen et al. 2019, Martin et al. 2022), and to envisage diverse forms of spatio-temporal environmental variation and/or change. However, for current purposes of demonstrating general conceptual principles, we do not explicitly invoke any specific real-life system. All parameters and notations are defined in Table 1.

**Table 1.**
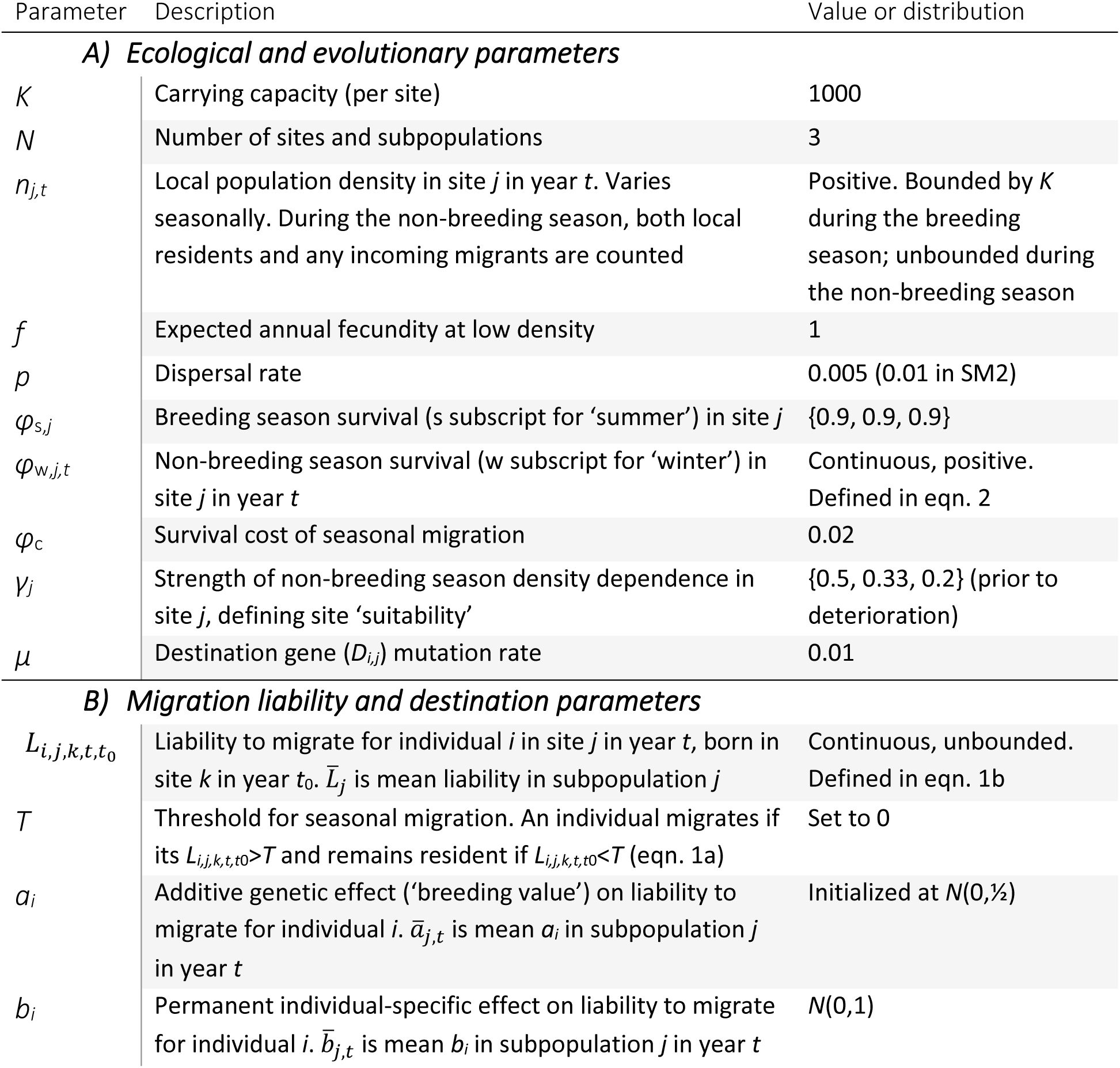
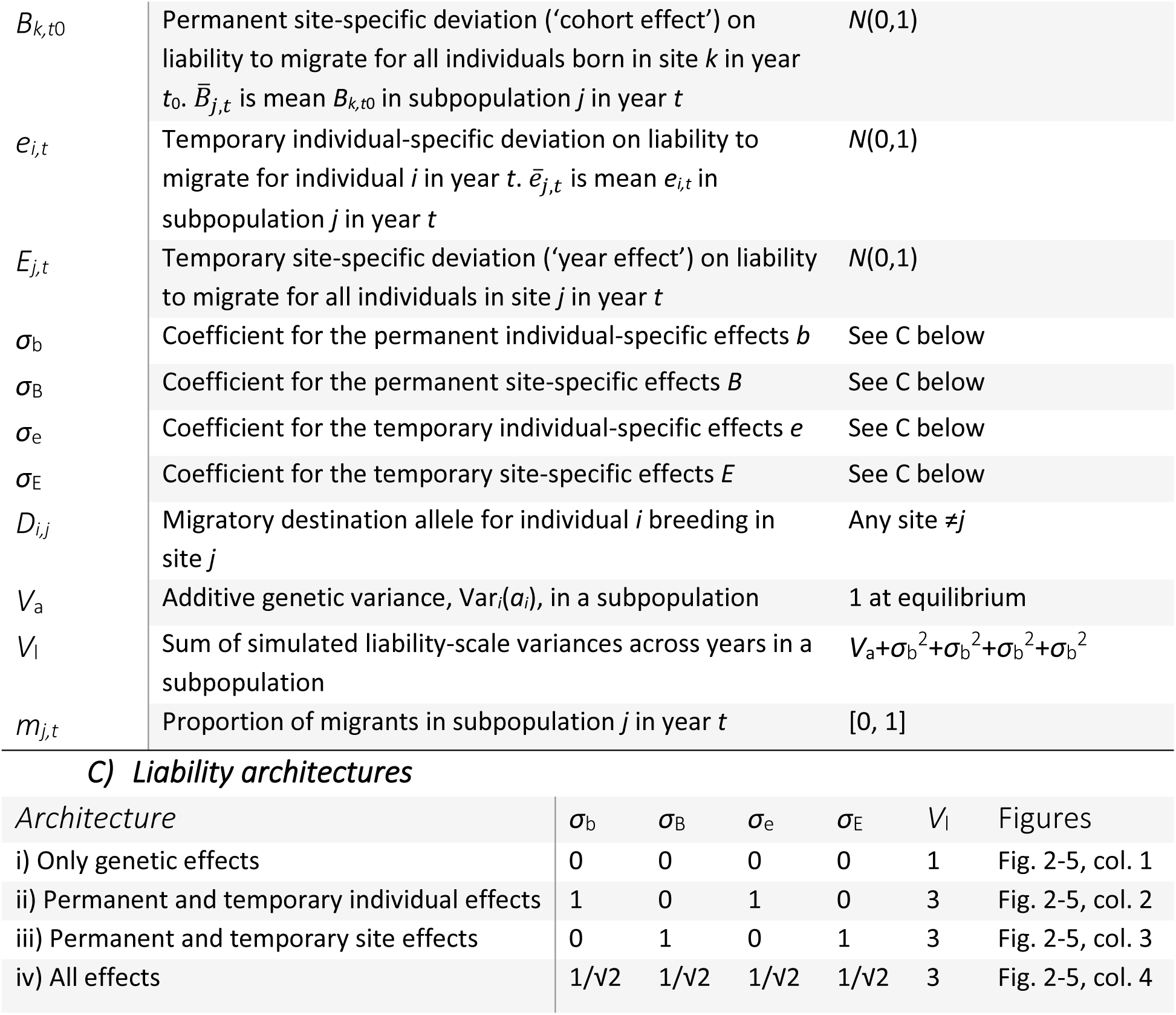
Summary of model parameters, notation, and simulated values: A) Ecological and evolutionary parameters. Bracketed values refer to sites A, B and C respectively. B) Migration liability and destination parameters. C) Parameterizations and summaries of four focal liability architectures.

### Migration liability

We model seasonal migration versus residence as a quantitative genetic threshold trait (Fig. 1B,C), which is an empirically supported model for migratory polyphenism in birds (Pulido 2011), fish (Dodson et al. 2013), insects (Menz et al. 2019) and mammals (Berg et al. 2019), with clear links to statistical estimation (Acker and Reid 2025). An individual *i* born in site *k* in year *t*_0_, who survives to breed in site *j* in year *t*, has liability to migrate *L_i_*_,*j*,*k*,*t*,*t*0_ , which translates into seasonal migration (versus residence) if it exceeds a threshold *T* (taken as zero):

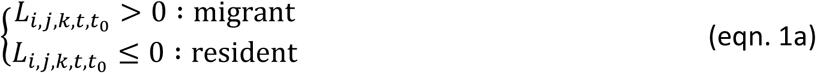

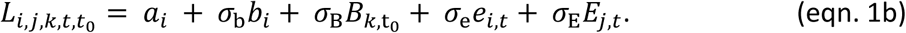

Here, *a_i_* is an individual’s additive genetic ‘breeding value’, which is inherited according to the infinitesimal model (Barton et al. 2017). Specifically, an individual’s value is drawn from a Gaussian distribution with mean equal to the mean *ai* of its parents and Mendelian sampling variance ½*V*_a_, where *V*_a_ is the additive genetic variance. This formulation assumes sexual biparental reproduction with no inbreeding, and implies that liability-scale genetic variation persists over time (interpretable as mutation-selection-drift-immigration balance).

Alongside *a_i_*, *L_i,j,k,t,t_*_0_ can also depend on four stochastic environmental components (eqn 1b, Fig. 1B,D). These include permanent individual effects *bi*, which are independently assigned to each individual at birth, representing developmental instability or irreversible plasticity in response to (micro-environmental) individual-specific early-life conditions. They also include permanent cohort effects *B_k,_*_t0_, which are shared by all individuals born in a given site *k* in year *t*_0_, representing developmental plasticity in response to (macro-environmental) site-specific early-life conditions. Further, they include temporary effects which vary among years, and which may also be individual-specific, *e_i,t_*, or site-specific, *E_j,t_*, representing labile (reversible) plasticity in response to micro-or macro-environmental variation respectively. Years *t* and *t*_0_ are identical for juveniles, and sites *j* and *k* are identical for individuals that do not disperse (see below).

Values of *bi*, *B_k,t_*_0_, *e_i,t_* and *E_j,t_* are drawn from normal distributions with mean 0 and variance 1 (i.e. interpretable as standardized ‘z-scores’), then scaled by parameters *σ*_b_, *σ*_B_, *σ*_e_ and *σ*_E_ respectively, which can be varied independently (Fig. 1D). Hence, we do not model plastic responses to explicit informative environmental cues, as often done when environmental change is postulated to affect cue reliability (Reed et al. 2010, Lande 2014, Scheiner et al. 2020, Clement et al. 2023). Rather, we examine the consequences of total environmental effects regardless of their multi-dimensional basis. This mirrors numerous empirical studies that estimate components of environmentally-induced variation without postulating their basis, which can stem from multiple (unmeasured) environmental variables (Kendall et al. 2011, Cam et al. 2016). The threshold trait formulation (eqn. 1a, Figs. 1C,S1) can then generate phenotypic plasticity non-additively (i.e. gene-by-environment interactions), where additive liability-scale environmental effects (i.e. liability-scale plasticity) might or might not cause phenotypic change depending on the proximity of the mean liability to the threshold (Reid and Acker 2022, Acker and Reid 2025). Our formulation of *L_i_*_,*j*,*k*,*t*,*t*0_ implies that migration does not depend on conditions at the destination, which are assumed unknown by individuals at the time of potential departure.

### Liability architectures

To examine spatio-seasonal eco-evolutionary dynamics arising given different environmental effects on *L_i_*_,*j*,*k*,*t*,*t*0_ in the context of wider environmental deterioration, we consider four focal sets of liability components (hereafter ‘architectures’), defined by setting the *σ*-parameters in eqn. 1b (Fig. 1B,D). First, as a baseline architecture for rapid evolutionary responses to environmental deterioration, we let migration liability *L_i_*_,*j*,*k*,*t*,*t*0_ be purely genetically determined, by setting all *σ*-parameters to zero, yielding liability-scale heritability *h*^2^=1. Second, we let permanent and temporary environmental effects on liability be purely individual-specific (i.e. *σ*_b_>0 and *σ*_e_>0), effectively capturing developmental and labile plasticity given some (implicit) individual-specific micro-environmental variation. Third, we let permanent and temporary environmental effects on liability be purely site-specific (i.e. *σ*_B_>0 and *σ*_E_>0). Here, *σ*_B_ generates a ‘cohort effect’ (i.e. a lasting phenotypic deviation shared by all individuals born in a certain site and year) and *σ*_E_ generates additional temporary (i.e. single-year) deviations, due to shared environmental conditions. Fourth, we consider an architecture with permanent and temporary individual-and site-specific effects on liability (all four *σ*-parameters >0), representing a more realistic case with multiple sources of trait variation.

In each case, we specify *V*_a_=1 and set the *σ*-parameters such that total simulated liability-scale environmental variance (*V*_a_+*σ*_b_^2^+*σ*_B_^2^+*σ*_e_^2^+*σ*_E_^2^) *V*_l_=3 for all architectures with environmental effects (Table 1C). Accordingly, *σ*-parameter values are 1 in the architectures with only individual-specific or only site-specific environmental effects, and 1/√2 in the architecture with all four effects. These values are chosen to illustrate general differences among liability architectures yielding broadly plausible heritability and repeatability. However, note that realized (site-by-year-specific) liability-scale heritabilities and repeatabilities are higher given site-specific effects, which generate less concurrent among-individual variance (and hence smaller *V*_l_ at a given time; see Results).

### Annual cycle

At the start of each breeding season, individuals in each subpopulation survive with site-specific probability *φ*_s,*j*_. Sexual reproduction then occurs. The total offspring produced in each site equals the smaller of the number of local breeders times expected fecundity *f*, and the site carrying capacity *K* minus current local breeding population size. This formulation effectively caps recruitment at *K*, allowing varying population sizes but not unlimited growth. Offspring are randomly assigned two parents from the local surviving adults. Adults are therefore potentially hermaphroditic, but this is inconsequential because no sex-specific effects are modeled.

After each breeding season, all individuals with liability *L_i_*_,*j*,*k*,*t*,*t*0_ >0 temporarily (i.e. seasonally) migrate to a different non-breeding season site from their breeding site *j*, while individuals with *L_i_*_,*j*,*k*,*t*,*t*0_ <0 remain resident at their breeding site (eqn. 1b). For seasonal migrants, destination is determined by a single haploid locus *D_i,j_*, with alternative alleles defining each possible destination site. An offspring’s *D_i,j_* allele is randomly inherited from one parent, and can mutate to an alternative allele (other than its breeding site *j*) with a small probability *μ*, allowing evolution of migratory destinations. Such large-effect loci determining migratory directions occur in nature (Sokolovskis et al. 2023), and higher *µ* could also be interpreted to represent greater environmental determination of migration destination. This simple architecture allows PMMPs to rapidly reach non-breeding season ideal free distributions (Haaland et al. 2026).

Density-dependent non-breeding season survival (or mortality) then occurs in each site. Here, an individual’s survival probability *φ*w,*j,t* decreases with current population density in site *j*, *n_j_*_,*t*_:

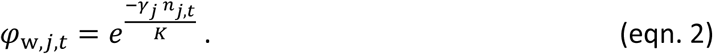

The site-specific parameter *γ_j_* adjusts the rate of decrease in survival with increasing density, representing seasonal habitat quality (higher *γ_j_* indicates steeper decrease, implying poorer quality). Mortality applies irrespective of whether an individual is resident at site *j*, or is an incoming seasonal migrant, and *nj*,*t* includes all individuals. It generates intrinsic frequency-dependent selection on expression of seasonal migration versus residence, and hence on underlying liability. This is because frequencies of seasonal movements affect *n_j_*_,*t*_ and hence *φ*_w,*j,t*_, while migrants and residents from a single breeding population by definition use different non-breeding season sites.

A separate density-independent mortality event then applies to seasonal migrants, who survive to return to their breeding site with probability *φ*c. This effectively imposes an overall cost of seasonal migration, including mortality during return movement, and any detriment of being an incomer at the non-breeding site. Finally, dispersal (i.e. changing breeding sites) occurs with a small, fixed probability *p*, generating some gene flow among breeding subpopulations. Dispersers are drawn independently of their genotype and any past migration decisions. If a disperser’s new breeding site *h* is the same as its seasonal migration destination, its *D* allele mutates to *D_i,h_,* ensuring that its migratory destination differs from its new breeding site.

To track spatio-seasonal population dynamics, the number of individuals present in each site is censused twice during each annual cycle: during the breeding season after mortality but before reproduction (pre-breeding census), and during the non-breeding season after density-dependent mortality, but before migration mortality and dispersal. The non-breeding season census includes all individuals present in a given site, encompassing residents and incoming seasonal migrants. It therefore represents the number of individuals that would be counted in a field census, not the number of individuals from the local breeding population that are currently alive.

### Spatial and ecological system

We embed the full annual cycle model within a landscape comprising *N*=3 sites (named A, B and C), with no explicit spatial arrangement (Fig. 1A). All sites are equally suitable for breeding populations (i.e. equal *φ*_s,*j*_), but non-breeding season suitability decreases from *j*=A to C (i.e., strength of density-dependence in non-breeding season survival, γ_w,*j*_, increases). This generates a simple PMMP, where subpopulations breeding in each site can evolve different liabilities to migrate and migratory destinations, driven by γ_w,*j*_ (Haaland et al. 2026).

Our parameterisations, including all directly specified and emerging survival probabilities (Table 1), generated expected adult lifespans of 4-5 years (2.5% of individuals live ≥10 years), broadly approximating life-histories of various partially-migratory birds, fish and mammals (Gillis et al. 2008, Berg et al. 2019, Morrison et al. 2021, Witczak et al. 2023). This life-history allows permanent individual-and/or site-specific effects on migration liability (representing forms of developmental plasticity) to persist across several years, and hence differ from among-year labile plasticity. Multi-year lifespans also allow individuals to express between-year phenotypic plasticity in seasonal migration versus residence.

### Environmental deterioration

For each liability architecture we then imposed gradual environmental deterioration starting in year 10050. For current conceptual purposes, we decreased non-breeding season suitability in site C, implemented as annually increasing density-dependence. Specifically, *γ*_C_ increased by 0.05/year from its starting point of 0.2, reaching 2.0 in year 10085, remaining at 2.0 subsequently (i.e., much higher than site A or B; Table 1). Such deterioration could represent, for example, gradually declining habitat availability or prey density.

To evaluate effects of the form of environmental deterioration, we ran three additional sets of simulations (Supplementary Material SM3). First, instead of gradually decreasing non-breeding season suitability in site C, we increased *γ*_C_ directly to 2.0 in year 10050. Second and third, to examine the degree to which results were caused by frequency-dependent selection on migration stemming from non-breeding season density-dependence, we enacted both gradual and sudden environmental deterioration in site C through fixed, density-independent mortality probabilities, rather than by increasing *γ*_C_.

### Effect of evolution

To isolate the effect of evolutionary change in *L_i,j,k,t,t_*_0_ on PMMP dynamics, we implemented control simulations where evolution during and following the period of environmental deterioration was prevented. For all offspring created after year 10050, genotypes (*a_i_* and *D_i,j_*) were drawn from the pre-deterioration population, not from the current parental generation. Thus, although the population’s genetic and phenotypic composition may still change somewhat due to differential survival of seasonal migrants versus residents within generations, no between-generation evolution occurs. These control simulations demonstrate evolutionary ‘rescue’ if population sizes following environmental deterioration were higher with evolution than without, and evolutionary ‘reduction’ if population sizes were lower with evolution than without. These uses of ‘rescue’ and ‘reduction’ do not necessarily imply that populations would respectively go extinct or remain fully unperturbed without evolution (i.e. representing “partial evolutionary rescue”, Schiffers et al. 2013).

### Simulation outputs and analyses

Starting from the initial equilibrium PMMPs for each liability architecture (Fig. 2A-D), we collected individual-level data comprising breeding and non-breeding season sites and liability *L_i_*_,*j*,*k*,*t*,*t*0_ and all its components for 50 years before imposing environmental deterioration, then for 150 years through the transition to the new stable environmental regime, for 20 independent replicate simulations.

**Figure 2:**
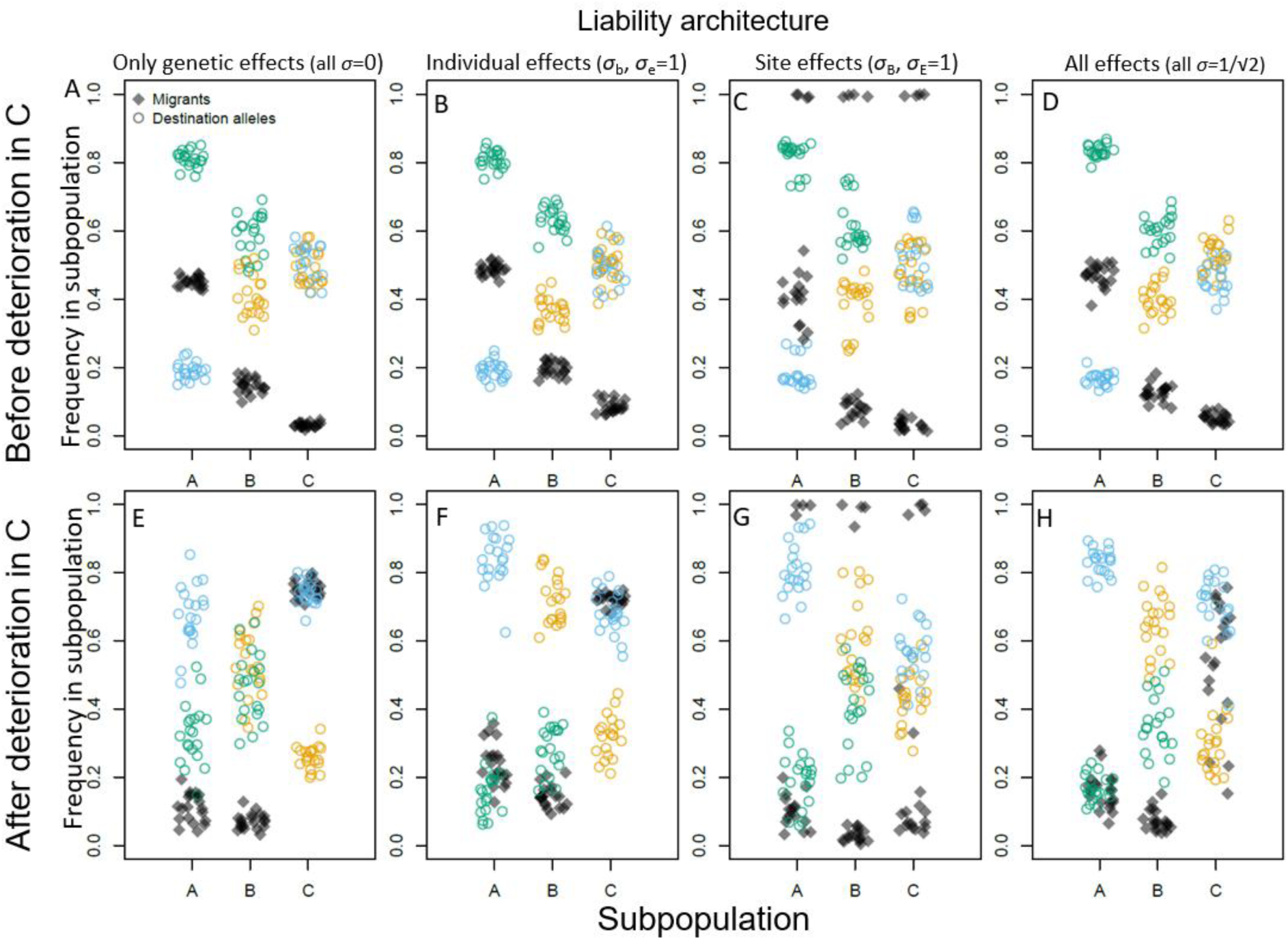
Overview of evolved partially-migratory metapopulation (PMMP) structures before (A-D), and after (E-H) deterioration of non-breeding season conditions in site C, given four focal liability architectures (columns). Graphs show mean frequencies of migrants (black diamonds) and *D_i,j_* alleles (i.e. migratory destination for individuals that migrate, denoted by orange (migrate to A), blue (migrate to B) and green (migrate to C) colored circles) in subpopulations A, B and C, averaged across (A-D) 20 observations from years 9050-10000 at 50 year intervals, or (E-H) the 20 final simulated years (10181-10200).

To quantify magnitudes of evolutionary change and strengths and impacts of selection on migration versus residence, and thereby reveal mechanisms causing observed evolutionary changes, we recorded between-year change in mean breeding value *a̅_j_*_,*t*_ − *a̅_j_*_,*t*−1_and proportions of migrants and residents *m* and 1–*m*, and computed non-breeding season survival probabilities *ϕ*_M_ and *ϕ*_R_ for migrants and residents (migrants with different destination *D_i,j_* alleles were pooled) and their mean Φ̅ = *φ*_M_*m* + *φ*_R_(1 − *m*) within each year, subpopulation and replicate. We then computed the selection differential (the difference in mean phenotype *m* after versus before selection) 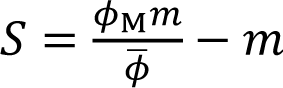 − *m*, and the selection gradient (the difference in relative fitness between migrants and residents) 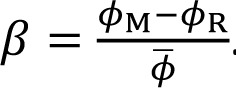 . These quantities are linked via the binomial phenotypic variance *m*(1-*m*), yielding *S*=*βm*(1–*m*).

To quantify realized heritability and repeatability we computed means and variances in breeding values, permanent effects and liabilities within each year, subpopulation and replicate (writing *b̅_j_*_,*t*_, *B̅_j_*_,*t*_ and *L̅_j_*_,*t*_ for the subpopulation-by-year-specific mean permanent individual deviations, permanent shared deviations and liabilities, respectively). Finally, to quantify dynamics of within-individual phenotypic plasticity in migration versus residence emerging from evolution of liabilities given temporary environmental effects, we classified individuals that lived ≥5 years as phenotypically plastic if they expressed both phenotypes at least once, and fixed (canalized) otherwise. We then computed the frequencies of plastic, fixed resident and fixed migrant individuals alive before, during and after evolutionary response to deterioration (Supplementary Material SM4).

All simulations were performed in R version 4.2.2 (R Core Team 2022). Code is available at http://datadryad.org/share/LINK_NOT_FOR_PUBLICATION/dKHkphLe2C2oilNju6jAkK_KsnXh4FT1t3UR0xNVwnc.

## Results

### Only genetic effects

Given the architecture with only genetic effects (i.e. *h*^2^=1; Figs. 3-5 left columns), non-breeding season environmental deterioration in site C caused dramatic spatio-seasonal eco-evolutionary dynamics spanning the PMMP, underpinned by evolution of liability for seasonal migration. Here, evolutionary responses diverge sharply between the subpopulation breeding in site C versus the other subpopulations (green versus orange and blue lines, Fig. 3). When non-breeding season suitability begins decreasing (Fig. 3, vertical dotted lines), there is immediate strong selection against residence in subpopulation C, and selection against seasonal migration to site C in the other two subpopulations. Consequently, there is a rapid evolutionary increase in *a̅*_C,*t*_, and corresponding evolutionary decreases in *a̅*_A,*t*_and *a̅*_B,*t*_(Fig. 3I, solid lines). Thus, initially ‘cryptic’ genetic variation allows subpopulation C to rapidly transition from almost fully resident into the most migratory subpopulation (Fig. 3A,E). Although the selection gradient in subpopulation C is steepest during early stages of adaptation when *a̅*_C,*t*_is very negative (Fig. 5I), the annual rate of evolution is fastest as *a̅*_C,*t*_ crosses the threshold (at zero), when phenotype frequencies are near 0.5 and the selection differential is greatest (Fig. 5A,E, grey shaded area). Then, *a̅*_C,*t*_ continues to increase, while *a̅*_A,*t*_and *a̅*_B,*t*_decrease, for several decades, with selection weakening and evolution slowing asymptotically as a new post-deterioration equilibrium is reached (Fig. 3A,E). Some among-replicate variation is also evident: replicates with more negative *a̅*_C,10050_ (i.e. at onset of environmental deterioration) take longer before *a̅*_C,*t*_ starts increasing (Fig. 3I), because genetic variation is too cryptic (i.e. centered too far from the threshold, and hence not phenotypically expressed and exposed to selection).

**Figure 3:**
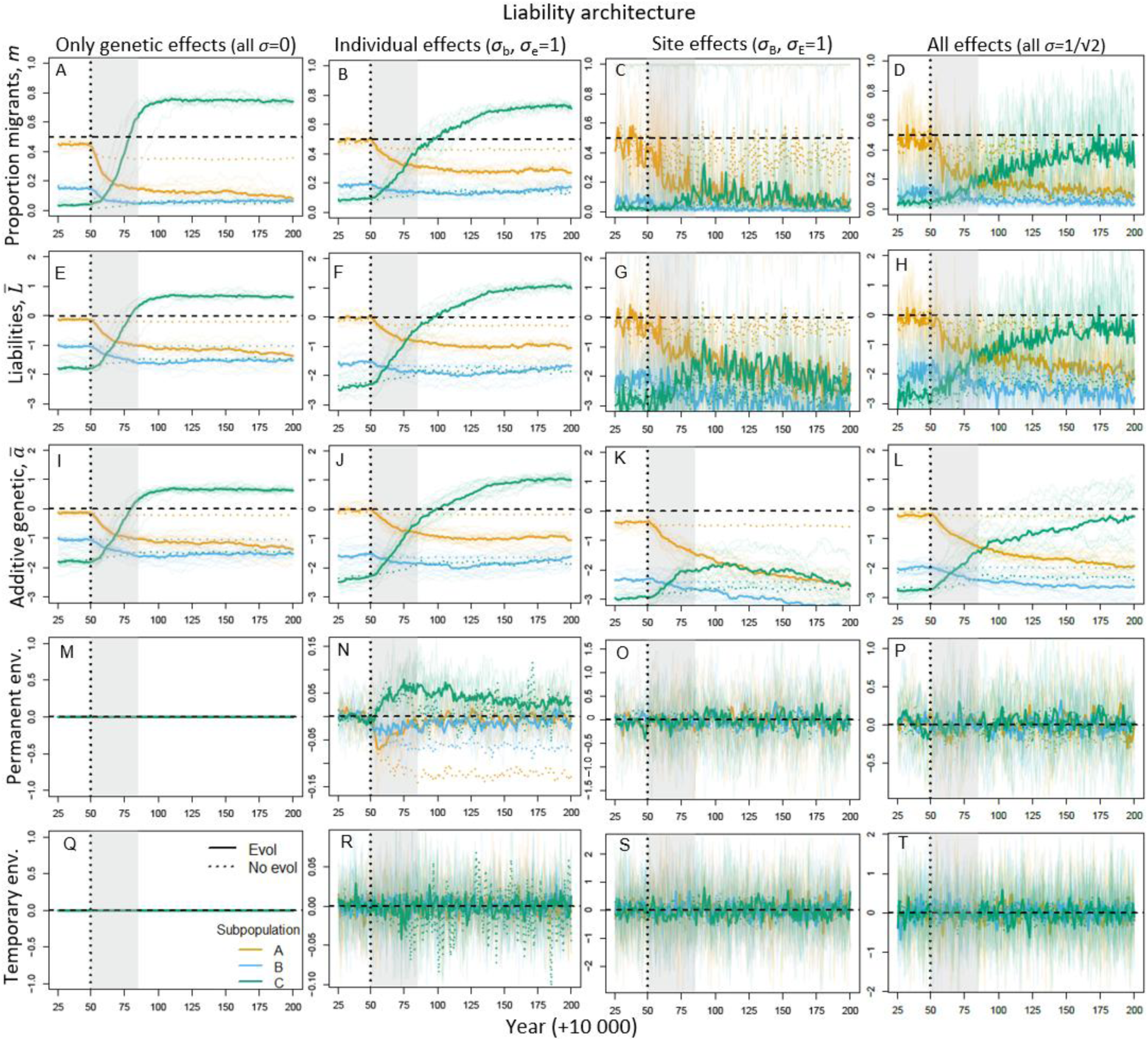
Evolutionary dynamics of seasonal migration during and following non-breeding season environmental deterioration in site C starting from year 10050 (dotted vertical line) and lasting for 35 years (grey shaded area). Columns show four liability architectures, comprising only genetic effects (column 1), permanent and temporary individual-specific environmental effects (column 2), permanent and temporary site-specific environmental effects (column 3), and all effects (column 4). Panels show trajectories for 150 years following onset of deterioration, for (A-D) proportion of migrants (*m*), (E-H) mean liabilities to migrate (*L̅*), (I-L) mean breeding values for liability (*a̅*), (M-P) mean permanent environmental deviations (*σ*_B_*B̅*+*σ*_b_*b̅*), and (Q-T) mean temporary environmental deviations (*σ*_E_*E_j,t_*+*σ*_e_*e̅*). In all panels, orange, blue and green denote subpopulations A, B and C respectively. Thick lines show medians across 20 replicate simulations and thin lines show individual realizations (20 in A-D, only 10 plotted in E-P for visual clarity). Horizontal dashed lines in A-H show the threshold value of zero, distinguishing liabilities that induce seasonal migration versus year-round residence, corresponding to 50:50 migration:residence (*m*=0.5) in I-L. Solid and dotted colored lines denote simulations with and without evolution respectively. In panels M and Q, all lines are on zero (with green printed last) because there are no environmental deviations given the liability architecture with only genetic effects. Note that y-axis ranges differ across columns in the two bottom rows.

These spatially structured evolutionary dynamics interact with spatial patterns of seasonal mortality and density-dependence to shape the emerging PMMP dynamics. Breeding season sizes of all three subpopulations rapidly decrease, especially in subpopulation C, as local residents (and incoming seasonal migrants) experience increased non-breeding season mortality (Fig. 4A, solid lines). Subpopulation C then starts to recover as it evolves to become more migratory and hence avoid the demographic cost of selection against residence. Here, replicates reaching smaller minimum population sizes correspond to those with more negative *a̅*_C,10050_ (correlation coefficient 0.78). Meanwhile subpopulations A and B evolve to become less migratory, and any migrants increasingly migrate to each other through evolution of *D_i,j_* (Fig. 2E). These processes increase non-breeding season densities in sites A and B (Fig. 4E, solid lines) and hence increase local non-breeding season mortality. Consequently, while the initial decreases in breeding season population sizes in subpopulations A and B are smaller than in C, these subpopulations continue to decrease as evolution progresses (Fig. 4A). Subpopulation sizes then return to being approximately equal, as the PMMP converges on its new ideal free non-breeding season distribution (Fig. 4A,E).

**Figure 4:**
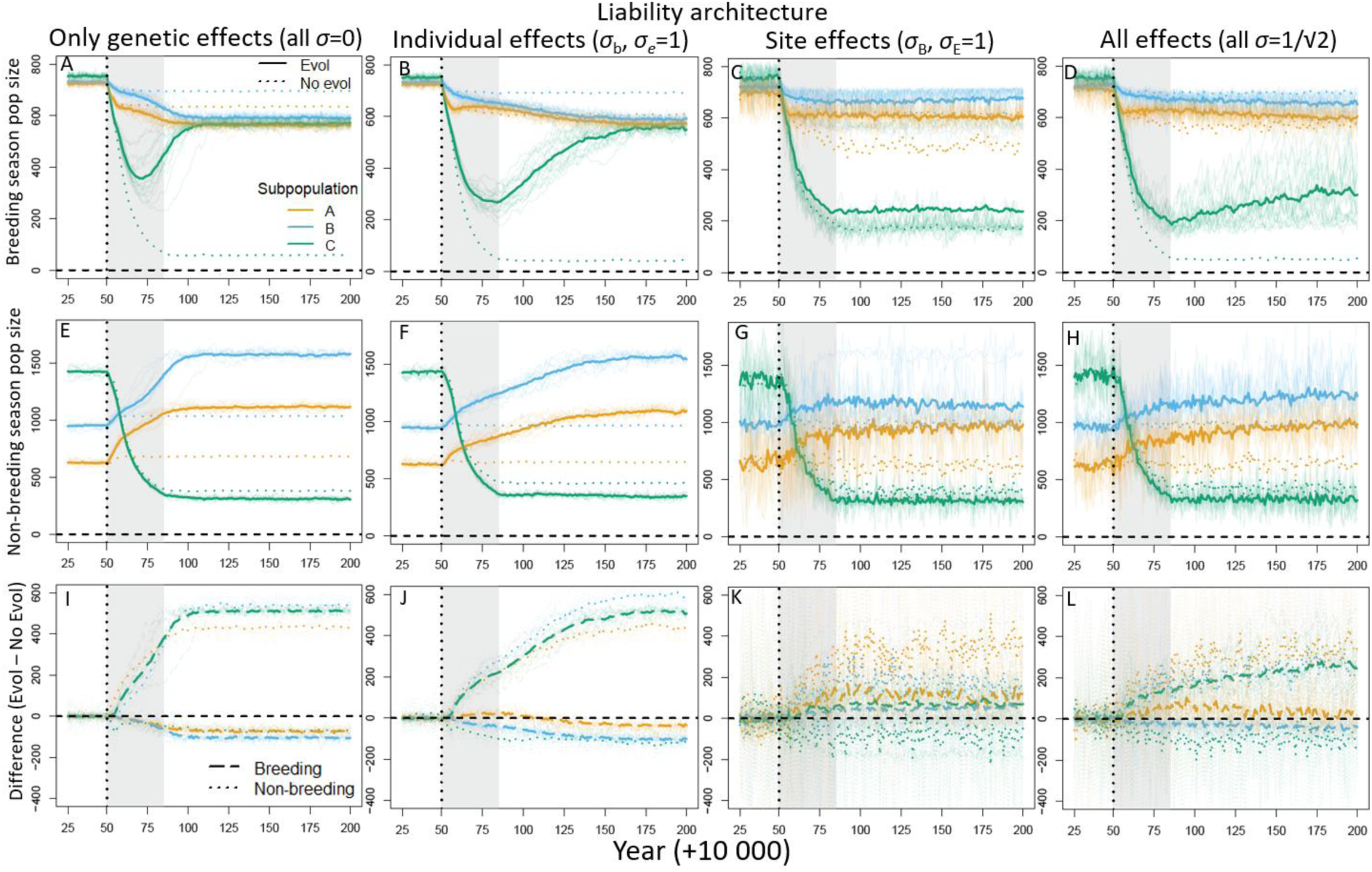
Spatio-seasonal population dynamics during and following non-breeding season environmental deterioration in site C starting from year 10050 (dotted vertical line) and lasting for 35 years (grey shaded area). Columns show four liability architectures, comprising only genetic effects (column 1), permanent and temporary individual-specific environmental effects (column 2), permanent and temporary site-specific effects (column 3), and all effects (column 4). Panels show trajectories over 150 years following onset of deterioration, for (A-D) breeding season adult population sizes, (E-H) non-breeding season population size for each site (i.e., number of individuals present, regardless of their subpopulation of origin), and (I-L) difference between local population sizes with vs without evolution (positive values indicate larger sizes with evolution than without). Orange, blue and green denote subpopulations A, B and C respectively. Thick lines show grand means across 20 replicate simulations and thin lines show individual realizations (20 in panels A-D, only 10 plotted in E-L for visual clarity). In panels A-H, solid and dotted lines denote simulations where evolution is respectively allowed and prevented. In panels I-L, dashed and dotted lines denote differences for respectively breeding and non-breeding season population sizes.

In comparison, in control simulations where evolution is prevented, breeding season sizes of subpopulations A and B stabilize soon after the environmental deterioration, while subpopulation C decreases to a new low equilibrium due to the high ongoing mortality of residents (Fig. 4A, dotted lines). The net effect of PMMP-wide evolution is therefore to increase total metapopulation size, by substantially increasing subpopulation C (local evolutionary ‘rescue’), but decreasing subpopulations A and B (local evolutionary ‘reduction’, Fig. 4I, dashed lines). Conversely, non-breeding season densities are lower in site C with evolution, but much higher in sites A and B (Fig. 4I, dotted lines). The PMMP structure therefore means that rapid evolutionary changes in *L_i,j,k,t,t_*_0_ following spatially-restricted environmental deterioration have notably diverging consequences for population densities across seasons and subpopulations.

### Individual-specific environmental effects

Adding permanent and temporary individual-specific deviations to *L_i,j,k,t,t_*_0_ (non-zero *σ*_b_ and *σ*_e_) clearly slows and reshapes evolutionary responses compared to those emerging given only genetic effects (Figs. 3-5, second column). It takes substantially longer for *a̅*_C,*t*_ to exceed zero (Fig. 3J; median across simulations 44 years since onset of environmental deterioration, versus 31 years given only genetic effects), while *a̅*_A,*t*_and *a̅*_B,*t*_ take longer to reach lower new equilibria. These differences are qualitatively expected, because part of the liability variance now stems from stochastic individual deviations that are not inherited (Table 1C, Fig. 5N). However, these deviations also serve to expose some genetic variation to selection when frequencies of migrants are initially low (*a̅*_C,*t*_ is far from the threshold), generating greater selection differentials during early stages of evolution (Fig. 5F, years 50-75) and hence faster evolution of *a̅*_C,*t*_ than might be expected simply given the lower heritability (*h*^2^≈^1^/_3_ vs 1). The deviations also cause transient increases then decreases in within-individual phenotypic plasticity, evident as *a̅*_C,*t*_evolves towards then away from the threshold (Fig. S11).

**Figure 5:**
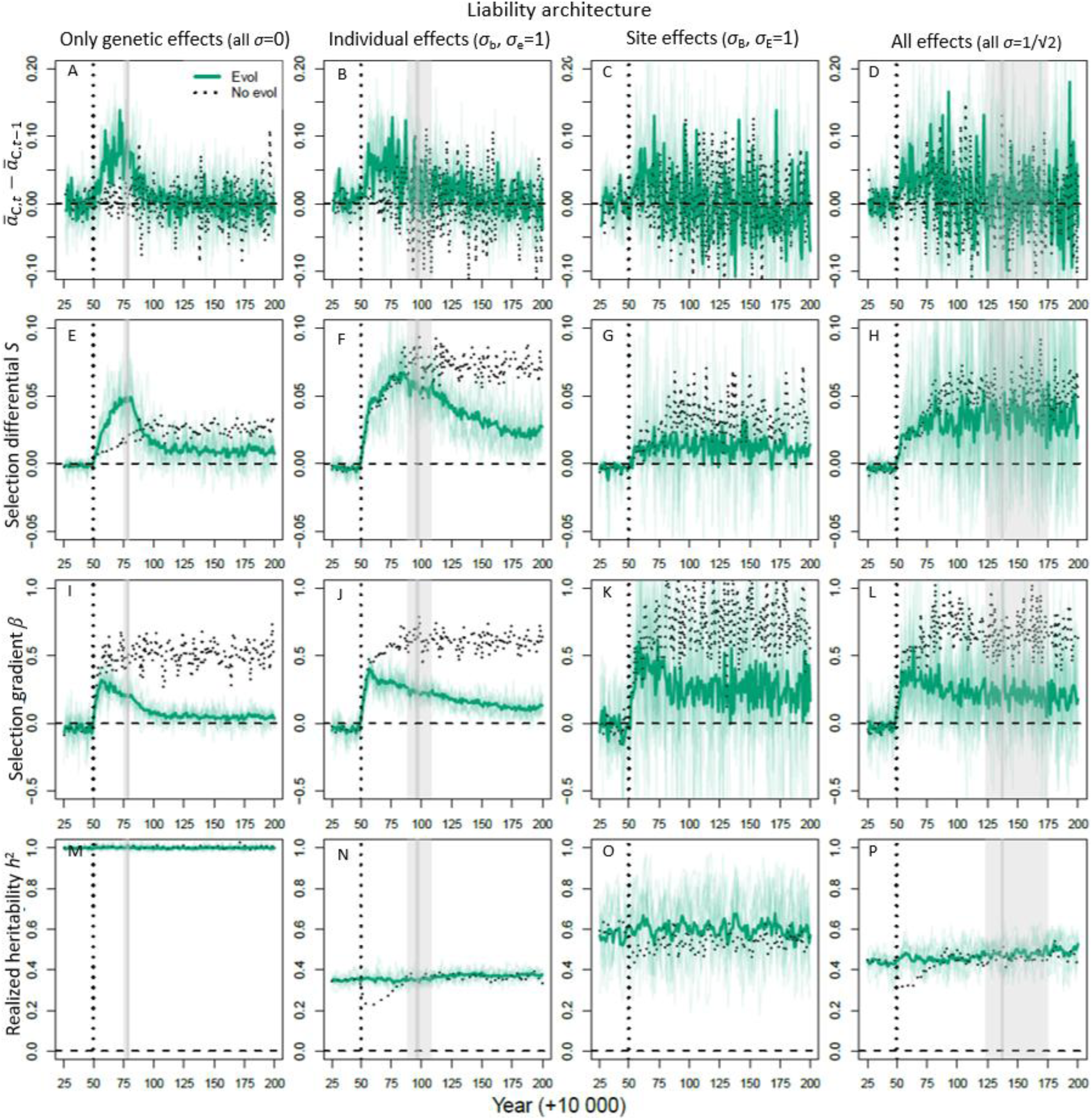
Trajectories of actual and potential evolutionary change following onset of environmental deterioration in site C (vertical dotted lines) across the four focal liability architectures (columns). (A-D) Between-year change in *a̅*_C,*t*_ in subpopulation C. (E-H) selection differentials *S* (difference between proportion of migrants after versus before non-breeding season survival selection). (I-L) Selection gradients *β* (difference in relative non-breeding season survival of migrants versus residents). (M-P) Realized liability-scale heritabilities *h*^2^=*V*_a_/*V*_l_. Grey vertical lines show the median years (across replicate simulations) at which *a̅*_C,*t*_ first exceeded the threshold of zero, and shaded grey bands show interquartile range (missing for the site effects architecture where no replicates evolved positive *a̅*_C,,*t*_). Thick green lines show grand means across 20 replicate simulations and thin lines show individual realizations. Dotted black lines show means in control simulations where evolution is prevented.

Additional shifts arise because selection on permanent individual deviations further increases *L̅*_C,*t*_, and decreases *L̅*_A,*t*_and *L̅*_B,*t*_(Fig. 3F,N). Here, selective removal of individuals whose environmental deviations make them year-round residents in subpopulation C, or seasonal migrants from subpopulations A and B, causes lasting deviations of *b̅*_A,*t*_, *b̅*_B,*t*_ and *b̅*_C,*t*_ from zero. This selection also reduces environmental variance in liability, thus increasing realized heritability slightly above the expectation of *h*^2^=^1^/_3_ (Fig. 5N). These population-level effects can persist until evolution is complete, through repeated selection on successive generations of individuals. Notably, *b̅*_C,*t*_ remains different from zero even a century after deterioration ended (Fig. 3N).

Because evolution is slower given individual-specific environmental effects than given only genetic effects, the recovery in breeding season size of subpopulation C, and hence the knock-on decreases in subpopulations A and B, are also slower (Fig. 4, second vs. first column). The new equilibrium PMMP structure (where subpopulation sizes return to being approximately equal) is only reached approximately a century after environmental deterioration ceased. Because of its protracted maladaptation, subpopulation C typically reaches a lower breeding season size before starting to recover (Fig. 4B vs. 4A). Non-breeding season densities in sites A and B also take much longer to increase (Fig. 4F vs. 4E), because fewer migrants arrive from subpopulation C due to its smaller size and slower evolutionary increase in migration.

### Site-specific environmental effects

Adding permanent and temporary site-specific deviations to *L_i,j,k,t,t_*_0_ (non-zero *σ*_B_ and *σ*_E_, Fig. 3-5, third column) causes notably different dynamics. Although the magnitudes of the environmental variance components are the same as in the architecture with individual-specific deviations, realized *h*^2^ is much higher (Fig. 5O) since shared site-specific deviations cause less among-individual environmental variance in liabilities. Yet, counter-intuitively, evolutionary increases in *a̅*_C,*t*_ upon environmental deterioration are considerably slower, and commonly cease or even reverse without crossing the threshold (at zero, Fig. 3G,K).

This failure of evolution is partly because the large stochastic among-year variation in *L̅_j_*_,*t*_in all subpopulations introduced by site-specific deviations causes among-year variation in non-breeding season densities. This often reverses the direction of selection on migration in subpopulation C due to density-dependent non-breeding season survival (Fig. 5G,K). Such fluctuating selection causes inherently asymmetric frequency-dependent evolutionary responses in threshold traits. For example, if a predominantly resident subpopulation C (with very negative *a̅*_C,*t*_) experiences selection for migration which increases the frequency of migrants towards 0.5 (and *a̅*_C,*t*_towards zero), a subsequent episode of similarly strong selection for residence will cause a greater evolutionary response, leaving the subpopulation even more resident (*a̅*_C,*t*_more negative) than it started. Thus, while selection is on average strong, reflected in greater selection gradients than given the other liability architectures (though more variable; Fig. 5K vs. 5I-J), responses to selection are much smaller and on average near zero (Fig. 5C). Further, unlike for permanent individual deviations, mean permanent site deviations *B̅_j_*_,*t*_do not show consistent responses to selection, but rather vary in sign among years (Fig. 3K). Repeated episodes of selective removal thus fail to directionally change *L̅_j_*_,*t*_, or hence adaptively alter the degree of migration.

Because evolution of *a̅*_C,*t*_is so slow, population density in site C decreases due to ongoing severe maladaptation (Fig. 4C). Consequently, because non-breeding season mortality is density-dependent, it gradually decreases, thereby weakening local selection for migration. Consequently, migration commonly never evolves to substantial proportions (Fig. 3C). Meanwhile, subpopulations A and B evolve even greater degrees of residence than given only genetic effects or individual-specific deviations on *L_i,j,k,t,t_*_0_ (Fig. 3A-C). Thus, site-specific deviations frequently cause a breakdown of the PMMP structure, where the three subpopulations are no longer interconnected by seasonal migration.

Consequently, breeding season sizes of subpopulation C never increase, and instead stabilize only slightly above those reached in control simulations where evolution was prevented (Fig. 4C,K). Meanwhile, subpopulations A and B are larger than without evolution (Fig. 4K), because they successfully evolve full residence and their non-breeding season densities (and resulting mortality) are not increased by incoming migrants from subpopulation C (Figs. 2G, 4G). Hence, given site-specific deviations, the population impacts of evolution are reversed compared with the previous two liability architectures (Fig. 4K vs. 4I,J).

These effects are further demonstrated through comparisons with additional simulations where environmental deterioration acts by decreasing absolute survival rather than by increasing density-dependence. Subpopulation C then more readily evolves migration because selection for migration no longer weakens as local population size decreases (Figs. S7-S10). Subpopulation C then increases in size, representing evolutionary rescue, while subpopulations A and B decrease. However, some replicates start off as fully migratory metapopulations (black diamonds at 1 in Fig. 2G). In these cases, neither form of environmental deterioration in site C greatly affects breeding or non-breeding season population sizes (Fig. 4C,G) or evolutionary trajectories (Fig. 3C).

To obtain initial stable PMMPs before applying environmental deterioration, we ran 20 replicate simulations with each of the four focal liability architectures (Table 1C) for 10000 years. Simulations were initiated with *a_i_*∼N(0,½) (implying 50:50 resident:migrant phenotypes and *V*_a_=1 at equilibrium), *D_i,j_* locus allele frequencies of ½ (i.e. equal probability of each alternative non-breeding site), and starting age Poisson distributed with mean 3. Evolutionary equilibria were quickly reached through evolution of *a̅_j_*_,*t*_(mean *a_i_* in subpopulation *j* year *t*) underlying *L_i_*_,*j*,*k*,*t*,*t*0_ , generating a partially migratory subpopulation A, predominantly resident subpopulation B, and near-fully resident subpopulation C (Fig. 2A-D, black diamonds). Evolution at the *D_i,j_* locus causes seasonal migrants from subpopulations A and B to mostly migrate to site C, whereas seasonal migrants from subpopulation C migrate to sites A and B approximately equally (Fig. 2A-D, colored circles), because the *D_i,j_* locus is almost neutral when migration is rarely expressed. These equilibrium PMMPs typically differed little between our focal liability architectures (Table 1C), although the site effects architecture sometimes causes evolution of fully migratory metapopulations (Fig. 2C, black diamonds at 1 in all three subpopulations). This is because of a stochastic positive feedback loop, where large among-year variation in frequencies of migrants and non-zero gene flow cause evolution of increasingly positive *a̅_j_*_,*t*_(Fig. S2,S3; Haaland et al. 2026). Overall, evolved PMMPs exhibited ‘ideal free’ distributions of non-breeding season site use, where annual mortality and thus breeding season subpopulation sizes were similar across sites (Haaland et al. 2026).

### All environmental effects

The liability architecture that includes all four individual-and site-specific environmental effects alongside additive genetic effects generally caused intermediate outcomes between the architectures with individual-specific or site-specific effects, where the extreme alternative outcomes resulting from purely site-specific effects are dampened. Notably, fully migratory metapopulations no longer occur (Fig. 2D vs. 2C), nor does subpopulation C so consistently fail to evolve migration from initial residence following deterioration (Fig. 3D vs 3C). Consequently, evolutionary rescue occurs more readily (Fig. 4D). These outcomes arise because, while the site-specific deviations still induce fluctuating selection (Fig. 5L) and thereby slow evolutionary responses, the individual-specific deviations expose cryptic genetic variation to selection when *a̅*_C,*t*_ is initially very negative, generating higher selection differentials (Fig. 5H). Initial evolution of increased *a̅*_C,*t*_and resulting increased proportions of migrants (Fig. 3D,L) thus increases local non-breeding season survival and resulting breeding season subpopulation size (Fig. 4D). Subsequent density-dependent non-breeding season selection for migration is therefore maintained over time (Fig. 5H vs. 5G), and *a̅*_C,*t*_evolves to cross the threshold before the simulation endpoint in approximately half the replicates (11/20, Fig. 3D). Yet, evolutionary responses are again stronger if environmental deterioration is density-independent (Supplementary Material SM3), with positive *a̅*_C,*t*_typically evolving (18/20 replicates), generating greater subpopulation sizes (i.e. improved evolutionary rescue). These outcomes emerge even though the consistent adaptive shifts in *b̅* evident given only individual-specific effects are obscured by the much larger stochastic site-specific deviations (i.e., Fig. 3P resembles Fig. 3O more than 3N). Yet, overall, site-specific effects on *L_i,j,k,t,t_*_0_ clearly fundamentally reshape spatio-seasonal eco-evolutionary dynamics, including generating diverse stochastic outcomes and greatly reduced evolutionary rescue (despite abundant underlying *V*_A_), remaining evident even long after environmental deterioration ceases (Figs. 3I-L, 4I-L). These conclusions remained similar given additional simulations where all four environmental variances equaled those in the architectures with only individual-or site-specific effects, thereby doubling the total environmental variance. They also remained similar when environmental deterioration was implemented suddenly rather than gradually, although speeds and magnitudes of responses can differ (Figs. S4-S5, S9-S10).

## Discussion

Environmental effects on focal traits, constituting plasticity, could facilitate or impede adaptive evolution and population persistence (Price et al. 2003, Chevin and Lande 2010, Ashander et al. 2016, Muñoz 2021). Yet, eco-evolutionary impacts of structurally different stochastic environmental effects are rarely considered, especially given traits with non-linear genotype-environment-phenotype relationships expressed within multi-year life-histories experiencing spatial and temporal environmental variation and change (Paenke et al. 2007, De Meester et al. 2019, Govaert et al. 2019, Klausmeier et al. 2020, Fouqueau and Polechová 2024, Khare et al. 2024, Urban et al. 2024). Our quantitative genetic model explicitly considers additive genetic effects and different forms of permanent and temporary individual-specific and site-specific environmental effects affecting the ecologically-critical trait of liability for seasonal migration embedded within a partially-migratory metapopulation (PMMP). Our simulations show how differing forms of stochastic environmental effects drive diverging evolutionary, phenotypic and population dynamic responses to the same environmental deterioration, altering the rate and outcome of evolution and inducing contrasting instances of evolutionary ‘rescue’ and ‘reduction’ across subpopulations. We thereby demonstrate how diverse forms of plasticity, non-linear trait architectures and density-dependence in fitness, all of which are commonplace in nature, can interact to generate divergent and counter-intuitive eco-evolutionary outcomes.

### Evolutionary dynamics of dichotomous threshold traits

Even given constant underlying additive genetic variance (*V*_a_, and hence fundamental evolutionary potential) across all simulations, emerging rates and outcomes of liability-scale and phenotypic evolution varied substantially, depending on whether environmental deviations were individual-specific or site-specific. While overall evolutionary responses to environmental deterioration were slower given individual-specific deviations than given only additive genetic effects, initial evolution was not slowed as much as might be simply expected given the substantially reduced liability-scale heritability. This is because the non-linear genotype-environment-phenotype relationship underlying dichotomous phenotypic traits causes frequency-dependent responses to selection, which can be facilitated by environmental deviations.

Specifically, environmentally-induced among-individual variation in liability can push phenotype frequencies (in our case, frequency of migration) towards 0.5 even if the mean environmental deviation is zero, increasing evolutionary responses and exposing otherwise cryptic genetic variation to selection when mean liability (*L̅_j_*_,*t*_) is far from the threshold. Selection differentials consequently become greater than given purely additive genetic effects on liability, despite similar selection gradients (relative fitness of migrants versus residents), especially following the onset of deterioration. This effect is also found in one of the few previous models investigating the role of trait architecture on the evolution of partial migration (de Zoeten and Pulido 2020), where individual-specific environmental deviations accelerated adaptive evolution of residence when migration was formulated as a discrete threshold trait, but not when it was continuous.

These phenomena are further exacerbated by the effects of selection on permanent individual deviations, where multi-year life-histories and repeated episodes of selective disappearance cause lasting non-heritable shifts in *L̅_j_*_,*t*_and hence in phenotype frequencies, further affecting evolutionary responses. Selective disappearance of consistently differing individuals is well known to affect population mean phenotypes, for example causing cross-sectional age-specific changes (Cam et al. 2002, van de Pol and Verhulst 2006, Wynn et al. 2025). Yet, this process is surprisingly rarely considered in the context of eco-evolutionary dynamics (Forsythe et al. 2024). Given our current parameterisations with equal or smaller permanent individual variance than *V*_a_, the impact of selective disappearance on *L̅_j_*_,*t*_is small compared to that of evolution. However, permanent individual variance may often greatly exceed *V*_a_ in nature, including in liability for seasonal migration (Gillis et al. 2008, Arnaud et al. 2013, Acker et al. 2023). Given that adult lifespans of many partially-migratory species are moderately long (e.g. seabirds (Acker et al. 2023), ungulates (Berg et al. 2019), marine mammals (Geijer et al. 2016)), selection on permanent individual deviations could substantially shape population and evolutionary responses to changing seasonal environments. Indeed, in partially-migratory European shags (*Gulosus aristotelis*), episodes of storm-induced selection on permanent non-heritable deviations substantially affected population mean liability to migrate and phenotype frequencies in subsequent years (Acker et al. 2023).

However, specifying site-specific rather than individual-specific environmental deviations on liabilities caused very different evolutionary dynamics. Contrary to simple expectation given the reduced among-individual environmental variation and resulting higher liability-scale *h*^2^, evolutionary responses were notably slower than given individual-specific deviations, to the degree that seemingly adaptive evolutionary shifts were never achieved. These outcomes can again be explained by the consequences of the non-linear genotype-environment-phenotype relationship. Here, site-specific deviations generate large among-year variation in frequencies of residents and migrants, on average pushing frequencies away from 0.5 when *L̅_j_*_,*t*_is near the threshold. This weakens the frequency-dependent responses to selection, particularly impeding adaptation when the novel environmental conditions require evolution across the threshold (as in our subpopulation C). Selection differentials are thus greatly reduced, despite larger (although more variable) selection gradients than given individual-specific environmental deviations.

Furthermore, in the context of PMMPs, resulting uncorrelated among-year variation in the frequencies of migrants and residents in all subpopulations generates considerable among-year spatial variation in non-breeding season densities. This changes the relative fitnesses of migrants and residents from any one subpopulation through density-dependent non-breeding season survival, causing frequent reversals in directions of selection. Such reversals can cause asymmetrical and constrained evolutionary trajectories because of intrinsic frequency-dependence in evolutionary responses. For example, if one episode of selection increases the frequency of initially rare migration towards 0.5, then an episode of equally strong selection for residence the following year will lead to even less migration than the starting point, thus impeding net evolution across the threshold. Our simulations show that, together, these mechanisms can be strong enough to prolong maladaptation sufficiently to substantially reduce the size of subpopulation C. This in turn alleviates density-dependent selection for migration, producing alternative eco-evolutionary outcomes that ultimately erode migratory connectivity and eradicate the interlinked PMMP structure.

These impacts of individual-specific versus shared site-and/or cohort-specific environmental deviations on rates of evolution are intrinsic to traits with non-linear genotype-environment-phenotype relationships, and hence will apply far beyond the context of seasonal migration versus residence (Roff 1996). Yet, they have scarcely been considered in existing theory on eco-evolutionary responses to environmental change, which typically focusses on traits that are continuously distributed on observed phenotypic scales with linear genotype-environment-phenotype relationships and simple forms of plasticity (Chevin et al. 2010, Tufto 2015, Lyberger et al. 2021, Clement et al. 2023). Such constructions imply that evolutionary responses are unbounded, bidirectionally symmetrical and independent of the trait mean (and hence frequency-independent), which is implausible for numerous life-history traits that proximately shape population dynamics.

Yet, in studies of diverse traits in wild populations, quantitative variation (on phenotypic or latent scales) is commonly partitioned into additive genetic and permanent and temporary individual and year variances, and sometimes further into non-additive genetic, spatial and/or cohort variances, often revealing substantial components (Wilson et al. 2010, Charmantier et al. 2014, Hamel et al. 2018). Our simulated permanent and temporary site-specific deviations, which critically affected eco-evolutionary outcomes, would appear as cohort and year variances respectively, providing direct links with empirical observations. In contrast, theory on evolutionary rescue and eco-evolutionary dynamics commonly envisages organisms with simple life-histories, with a single selective event per discrete generation, eliminating the distinction between developmental and labile plasticity and eradicating any multi-year processes (Gomulkiewicz and Holt 1995, Lande 2009, Chevin et al. 2010, Orr and Unckless 2014, Klausmeier et al. 2020, Lyberger et al. 2021, Clement et al. 2023, Xu et al. 2023). Meanwhile, models that include more complex life-cycles and overlapping generations typically do not consider complex structures of environmental effects on vital rates (Barfield et al. 2011, Cotto et al. 2019). Our model, which considers diverse forms of environmental effects underlying a dichotomous quantitative genetic trait within an iteroparous (meta)population with overlapping cohorts, therefore reveals how relaxing typical assumptions to better capture environmentally-induced evolutionary demography can generate qualitatively different eco-evolutionary dynamics.

### Spatio-seasonal metapopulation dynamics

PMMPs, comprising reproductive subpopulations interlinked by non-breeding season migrants, are commonplace across diverse taxa (e.g. Papastamatiou et al. 2013, Allen et al. 2019, Arnekleiv et al. 2022, Payo-Payo et al. 2022), yet their eco-evolutionary responses to local environmental changes have scarcely been considered (Reid et al. 2018, Haaland et al. 2026). Our simulations capture internal and overall dynamics induced by spatially and seasonally restricted environmental deterioration, revealing strong divergent evolutionary, phenotypic and population dynamic responses across subpopulations stemming from evolving seasonal migration and resulting degrees of non-breeding season sympatry. For example, adaptive evolutionary increases in migration in the subpopulation most directly impacted by the seasonal environmental deterioration (here, subpopulation C) cause higher non-breeding season densities elsewhere, and hence reduce annual survival and thus breeding season densities in those subpopulations compared with control simulations where evolution was prevented. Hence, local adaptive evolution causing evolutionary rescue in one subpopulation simultaneously reduces resulting subpopulation sizes elsewhere. Existing evolutionary rescue models do not accommodate such spatio-seasonal eco-evolutionary dynamics because they consider a single breeding population experiencing seasonality (Kaitala et al. 1993, Griswold et al. 2010, Shaw and Levin 2013, Vélez-Espino et al. 2013, Ohms et al. 2019), or consider multiple subpopulations connected by permanent movements (i.e. dispersal) without seasonality (e.g. Schiffers et al. 2013, Bourne et al. 2014). In contrast, our PMMP model delivers on previous calls for evolutionary rescue theory to include ecologically richer contexts and non-linear genotype-environment-phenotype maps (Klausmeier et al. 2020). Incorporating this complexity further reveals that eco-evolutionary responses to sustained environmental deterioration, or stochastic extreme events (Haaland et al. 2026), can result in very long-lasting and spatially-widespread transient dynamics (even in simple cases with high heritability), indicating that PMMPs may rarely be at equilibrium in nature.

The potential for negative effects of adaptive evolution on population size or persistence, involving increased relative fitness but decreased absolute fitness, has been shown in other contexts. For example, evolution of extravagant secondary sexual traits that increase mating success can decrease survival and hence the growth rate of single populations (Rankin et al. 2007, Kokko 2011). Evolutionary rescue models have also shown how reduced adaptive potential can facilitate population persistence, either because increased genetic variation (despite providing the ‘fuel for adaptive evolution’) also contributes to variance load in a population tracking a moving environmental optimum (Lynch and Lande 1993, Klausmeier et al. 2020), or because faster adaptive evolution during extreme one-off events given higher heritability may be maladaptive when the event ends (Lyberger et al. 2021). In contrast, we demonstrate how adaptive evolution can drive subpopulation declines due to eco-evolutionary dynamics emerging in density-regulated metapopulations interlinked through evolving seasonal migration.

### Implications and extensions

Our current parameterizations and simulations are designed to highlight key conceptual points, not to predict realistic eco-evolutionary dynamics of any particular system. Simulating different spatio-temporal structures, parameter values and forms of environmental change would yield quantitatively, and perhaps qualitatively, different specific outcomes. Future ambitions should therefore be to quantify impacts of different forms and magnitudes of environmental effects under diverse hypothetical or real regimes of environmental change, including rates, directions and spatial patterns of environmental deterioration and/or sequences of stochastic extreme events that could cause fluctuating rather than directional selection (Baeckens and Donihue 2025, Haaland et al. 2026). To facilitate this, our quantitative genetic model is constructed to allow direct parameterization from statistical quantitative genetic analyses applied to individual-based data collected in wild populations (Kruuk et al. 2014). This differs from most previous evolutionary theory on dichotomous phenotypes such as seasonal migration versus residence, which employs genetically implicit approaches that are less empirically tractable (e.g. evaluations of evolutionary stable phenotypic states; Kaitala et al. 1993, Kokko 2011, Shaw et al. 2019).

Our current simulations consider stochastic environmental deviations on liabilities to migrate, which could also be interpreted as resulting from latent reaction norms of plasticity on stochastic environmental variation (Supplementary Material SM1). Yet, differing eco-evolutionary dynamics could occur given liability-scale plasticity in response to informative cues that effectively predict fitness outcomes (i.e. directly adaptive plasticity). Indeed, one focus of wider theory on plasticity and evolutionary rescue concerns the degree to which cues reliably predict environments across successive timepoints (Lande 2014, Ashander et al. 2016, Clement et al. 2023, Martin et al. 2023). However, such models typically envisage populations in single locations with no seasonality, where traits both respond to and affect fitness through some local temporally-autocorrelated environmental variable. Yet, such adaptive plastic adjustment of seasonal migration is famously challenging (Bauer et al. 2020), because breeding season conditions may not predict non-breeding season conditions either locally or elsewhere. One exception may be seasonal population density, which may provide some information on likely densities in the upcoming season, also conditional on the liabilities (and hence migration) of all other subpopulation members. Modelling, and empirically estimating, permanent and/or temporary effects of local density on *L_i_*_,*j*,*k*,*t*,*t*0_ could therefore be a useful next step towards characterising spatio-seasonal eco-evolutionary dynamics under environmental change.

## Supplementary materials

for “Differing components of plasticity in threshold traits cause diverging eco-evolutionary responses to spatio-seasonal environmental deterioration”

### SM 1: Liability-scale and phenotypic plasticity in threshold traits

Here we illustrate how additive genetic and environmental effects on the liability *L* underlying threshold traits, and on resulting phenotypic expression, can be envisaged to stem from a reaction norm on the liability scale (Fig. S1A; see also Reid and Acker 2022). Specifically, the liability consists of an additive genetic effect and a set of one or more environmental components ***σ*_x_*x*** (bold characters representing vectors), where each *x*∼N(0,1) is the standardized effect (*z*-score) of one type of stochastic environmental variable, and each *σ*_x_ scales the magnitude of the effect (Table 1). Our current model and simulations consider permanent individual-specific effects *b*, permanent site-specific effects *B*, temporary individual-specific effects *e* and temporary site-specific effects *E*. The standardized effect of the average environmental conditions (*x*=0) then represents the individual’s breeding value *a_i_*. This can be seen as the mean-intercept of the liability-scale reaction norm, and the scaling parameter *σ*_x_ for a specific environmental component *x* can be seen as the slope of the liability-scale reaction norm with respect to the given environmental variable (eqn. 1b). Since these environmental conditions are implicit rather than explicit in our model, with liability effects corresponding to stochastic environmental draws rather than informative cues about current (or future) conditions, this approach can also be seen as a character-state approach which does not necessarily imply a linear reaction norm (Via et al. 1995). Then, this underlying liability *L*=*a_i_*+Σ*_x_*{*xσ_x_*} translates into alternative phenotypes of seasonal migration or residence depending on whether or not *L* exceeds the threshold value of 0 (eqn. 1a; dashed horizontal line in Fig. S1).

Further, given non-zero *σ*_e_ or *σ*_E_ (reaction norm slopes in response to temporary environmental components), an individual *i*’s liability can change between timepoints *t* as environmental conditions specific to individual *i, e_i,t_*, or to site *j*, *E_j,t_*, change. Depending on the distance of an individual’s mean liability *a_i_* from the threshold, these changes may translate to varying degrees into phenotypic change (switching from resident to migrant or *vice versa* from one year to the next). Fig. S1B shows examples of differences among individuals in such among-year phenotypic-scale plasticity (i.e. expression of migration vs. residence) versus canalization emerging due to phenotypic GxE, despite identical liability-scale plasticity. Individuals with intercepts closer to the threshold (e.g. middle reaction norm), express phenotypic plasticity even under moderate deviations from mean environmental conditions, whereas individuals with intercepts further from the threshold (e.g. darkest and lightest blue reaction norms) only do so given sufficiently large environmental deviations. Accordingly, the threshold trait formulation allows expressing both plasticity as well as “fixed” (or “obligate”) residence or migration, although seemingly obligate phenotypes are in fact not constrained to that phenotype. Note also that considerable ‘cryptic’ variation in *a_i_, σ* and *L* values could exist in highly canalized individuals with *a_i_* far from the threshold, since such liability-scale variation does not translate into phenotypic variation under typical environmental conditions. However, as our environmental deterioration simulations show, such variation may be expressed and exposed to selection when the environmental regime changes, shaping eco-evolutionary responses (Fig. 3).

**Figure S1:**
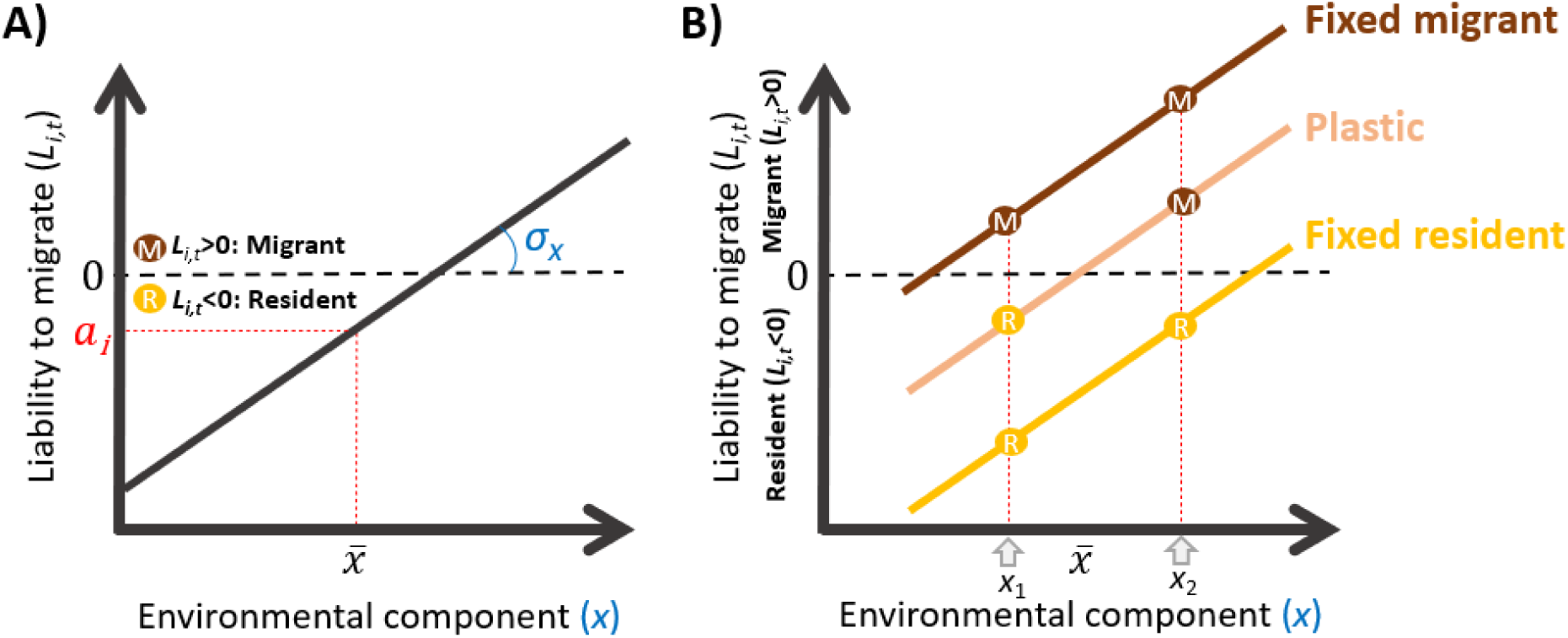
Examples of the threshold trait construction with liability-scale reaction norms and a dichotomous resulting phenotype, where the liability to migrate (y-axis) is shown as a function of one or more environmental components (x-axis). The threshold (black dashed line) is 0, such that individual *i* migrates in year *t* if its liability exceeds the threshold, and remains resident if not. The reaction norms can be envisaged as linear regressions, where the additive genetic effect *a_i_* on liability represents the elevation, and the *σ_x_* parameters represent the slope of the reaction norm with respect to environmental variable *x* (but, given that environments are implicit, reaction norms are not necessarily linear). A) Reaction norm for an individual which is on average resident (*ai*<0), but may migrate when the environmental deviation *x* is sufficiently positive, thus expressing among-year phenotypic plasticity. B) Reaction norms for three individuals (colours) with different *a_i_* (elevations), with phenotypes (migrant “M”, brown circles; resident “R”, yellow circles) evaluated at two example time points with environmental conditions *x*_1_ and *x*_2_, illustrating emergent among-individual differences in phenotypic plasticity despite identical liability-scale plasticity *σx*, due to GxE that emerges on the phenotypic scale. Cryptic genetic variation is also present in given environments, e.g. yellow and pink individuals both express the same phenotype in *x*_1_ (residence) despite having different *a_i_*, which only manifests phenotypically in environmental condition *x*_2_. Note that given different realizations of environmental conditions, the classifications of individuals (“fixed migrant”, “plastic”, “fixed resident”) shown here could change.

### SM 2: Evolved differences in partially migratory metapopulations among liability architectures

In the simulations run to generate initial equilibrium PMMPs (to which long-term environmental deterioration was then applied), we noted that liability architectures with larger stochastic deviations to *L_i_*_,*j*,*k*,*t*,*t*0_ (i.e. larger values of *σ*-parameters, eqn. 1b) tend to evolve increased degrees of seasonal migration (versus residence, Fig. S2). This occurs for both permanent and temporary environmental effects, but increases tend to be larger for site-specific than for individual-specific effects. Larger *σ* values, which are necessary to approach the low heritabilities and high individual repeatabilities seen in natural PMMPs, increase both mean and variance in levels of seasonal migration among simulation replicates. In some cases with substantial environmental stochasticity, an extreme of fully migratory metapopulations occurs. This outcome appears at lower *σ*-values given site-specific than individual-specific environmental deviations, as illustrated by the case when both have equal variances (within same rows in Fig. S2). This maladaptive outcome is in part due to a positive feedback loop where gene flow from partially migratory subpopulations A and B into the fully resident subpopulation C causes increasing seasonal migration in C (Fig. S3), which increases non-breeding densities in sites A and B. This in turn selects for higher breeding values for migration in these subpopulations, such that continued gene flow makes subpopulation C even more seasonally migratory, strengthening the feedback loop (Haaland et al. 2026). In order to test this proposed mechanism we ran additional simulations with higher dispersal rates (*p*=0.01), in which fully migratory outcomes indeed occur more often and for lower levels of environmental stochasticity (compare Fig. 2B,C with Fig. S2C,D; and Fig S2A,B with Fig. S2E,F).

Accordingly, the starting points for the simulations of environmental deterioration differ slightly between our focal liability architectures (Fig. 2A-D), and for other conceivable architectures (Fig. S2). However, for most parameterizations that avoid fully migratory outcomes, these differences are minor compared to the changes that arise once the deterioration begins. Note that if we were to standardize pre-deterioration PMMPs among liability architectures by setting a given *a̅_j_*_,10000_for each subpopulation, rather than allowing these to emerge following evolutionary burn-in to equilibrium, we would risk confounding evolution in response to deterioration with evolution due to non-equilibrium dynamics of *a̅*.

**Figure S2:**
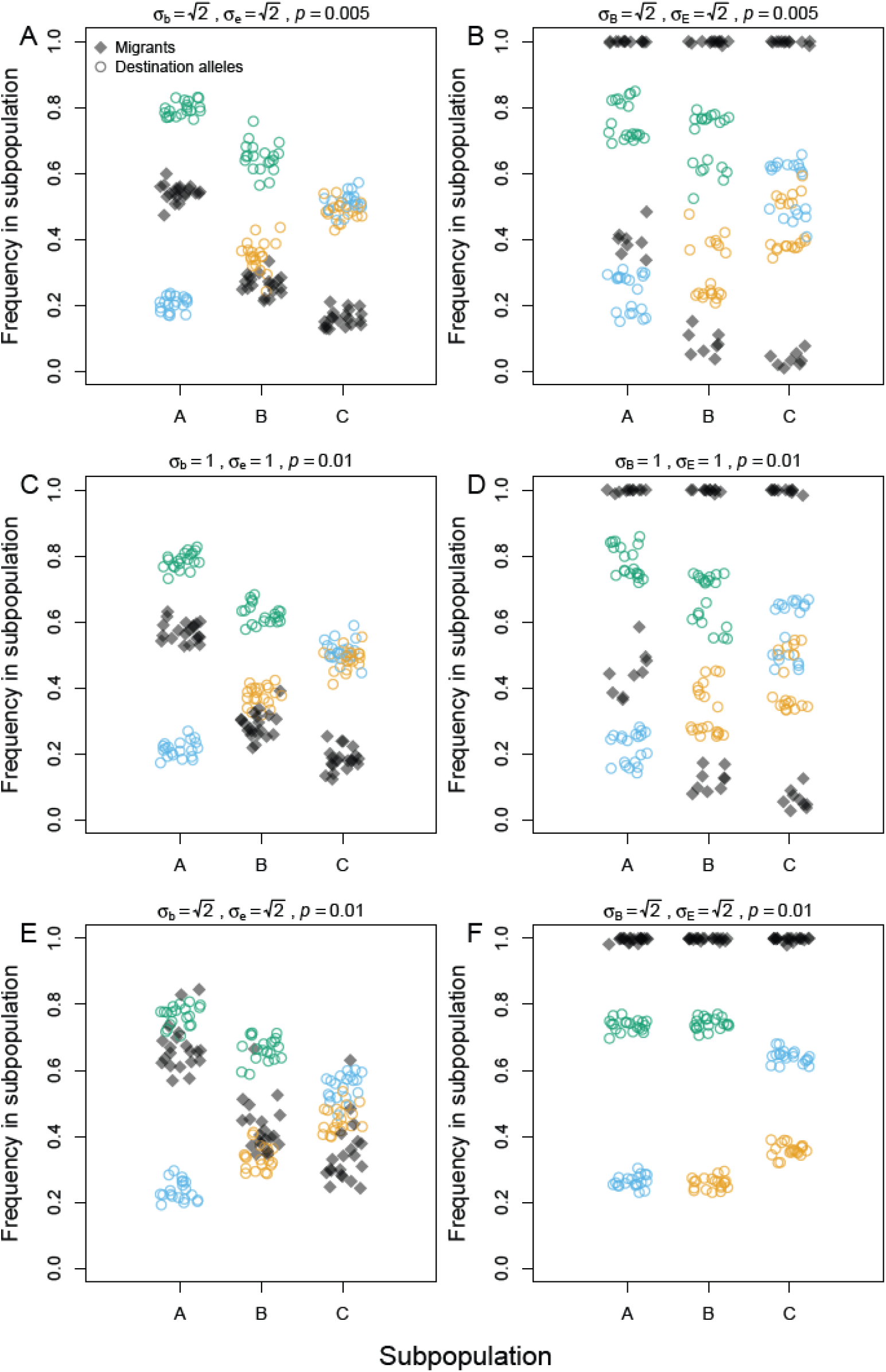
Effects of liability architectures and dispersal rate on evolved partially migratory metapopulations. Mean percentages of seasonal migrants (black diamonds) and *D_i,j_* alleles (colored rings; orange, blue and green respectively indicate migrating to site A, B or C) per subpopulation and replicate simulation, averaged over 20 observations at 50 year intervals between year 9000 and 10000. Permanent and temporary environmental deviations are individual-specific in panels A,C,E and site-specific in panels B,D,F. In panels A and B, dispersal rate *p*=0.005 (as in main text); in panels C-F *p*=0.01. Specified *σ*-values yield *V*l=5 in panels A,B,E,F, and 3 in panels C and D (as in main text). All *σ*-values not shown are set to 0. Other parameters are *γ*_n_ = {1/2, 1/3, 1/5}, *φ*_s_=0.9, *φ*_c_=0.02.

**Figure S3:**
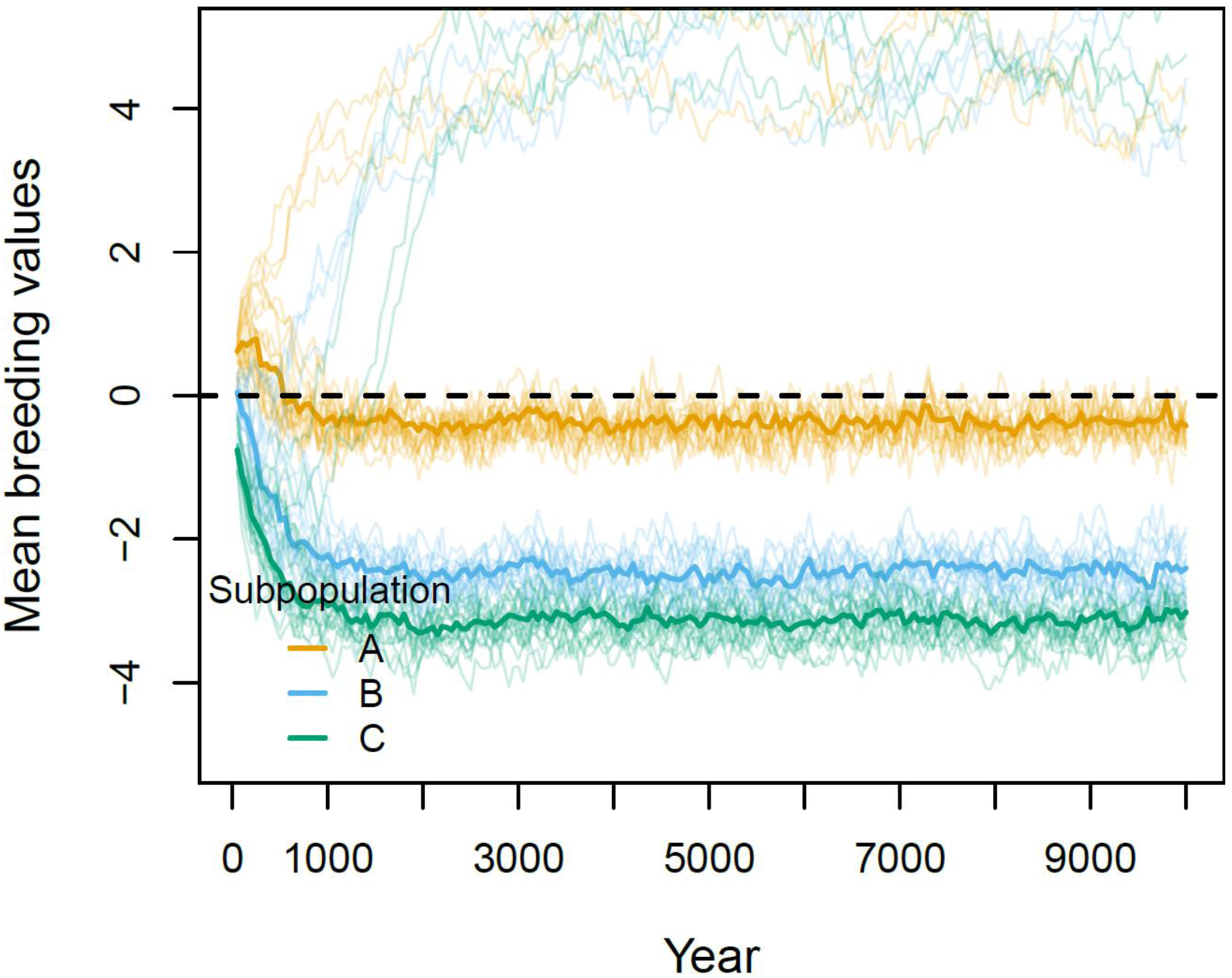
Pre-environmental deterioration evolutionary trajectories of mean breeding values (*a*) given the liability architecture with *σ*_B_=1 and *σ*_E_=1 (i.e. focal liability architecture ii, corresponding with Fig. 2C), illustrating the positive feedback loop occurring in a few replicates driving fully migratory metapopulations as an alternative evolutionary equilibrium. In replicates where subpopulation A starts off slightly more migratory (orange opaque lines), subpopulations B and C experience not only higher non-breeding season densities for residents (favoring migration), but also an influx through gene flow of higher (more positive) breeding values to subpopulations B and C (blue and green opaque lines). Both effects lead to evolution of ever more positive breeding values until all subpopulations are fully migratory, at which point selection on *a_i_* ceases.

### SM 3: Alternative formulations of environmental deterioration

We complemented the environmental deterioration scenario presented in the main text (gradual deterioration implemented by increases in density-dependent mortality) by considering three additional scenarios, creating a full factorial set of sudden or gradual deterioration in site C, and deterioration implemented via increases in density-dependent or density-independent mortality. The three additional scenarios complement our main conclusions regarding the roles of different environmental effects on migration liability in shaping responses to rapid deterioration, and their interactions with density-dependence.

Note that if environmental deterioration occurs somewhere else than in site C, the same general eco-evolutionary principles apply, although metapopulation-wide consequences of changes in non-breeding season conditions are greater when occurring in sites with higher non-breeding season densities (Haaland et al. 2026).

#### i) Sudden environmental deterioration, density-dependent

We first present the case where we let non-breeding season suitability in site C, formulated as strength of density-dependence, decrease suddenly rather than gradually. This scenario captures e.g. a sudden deterioration of habitat quality such as draining of a major wetland (Newton 2004). These two scenarios of gradual and sudden deterioration are commonly used in theoretical treatments of evolutionary rescue (Chevin and Lande 2010; Uecker et al. 2014; Ashander et al. 2016; Klausmeier et al. 2020). We set the strength of non-breeding season density-dependence *γ*_C_ to 2 directly in year 10050, i.e. the same as the end point of the gradual environmental deterioration scenario used in the main text.

For all liability architectures, onset of sudden environmental deterioration leads to an immediate population decline in all subpopulations that is much larger than the population decline occurring under gradual deterioration (Fig. S5A-D vs. Fig. 4A-D). As in the main text scenario, the regime shift selects for increased year-round residence in subpopulations A and B, and for increased seasonal migration in subpopulation C (Fig. S4A-D), but this immediate selection is now much stronger due to the extremely high mortality rate occurring in the first non-breeding season in site C when density is still at its pre-change optimum. However, following this initial decline, subpopulations A and B quickly return to near their pre-change population sizes (Fig. S5A-D), as they have become almost entirely resident after only a single year under the novel selection regime, thus avoiding the high non-breeding season mortality in site C (Fig. S4A-D).

When the liability architecture includes permanent and temporary individual-specific deviations (Figs. S4-S5, column 2), this shift to year-round residence is accelerated by a substantial decrease in mean permanent deviations *b̅_j_*_,*t*_ (about -0.15 and -0.05 in respectively subpopulations A and B, Fig. S4N orange and blue lines). These initial non-genetic responses are larger than in the gradual deterioration scenario, but also return to zero much faster, because the subpopulations sooner reach their post-deterioration optimal phenotypes (of near-full residence, Fig. S4B).

**Figure S4:**
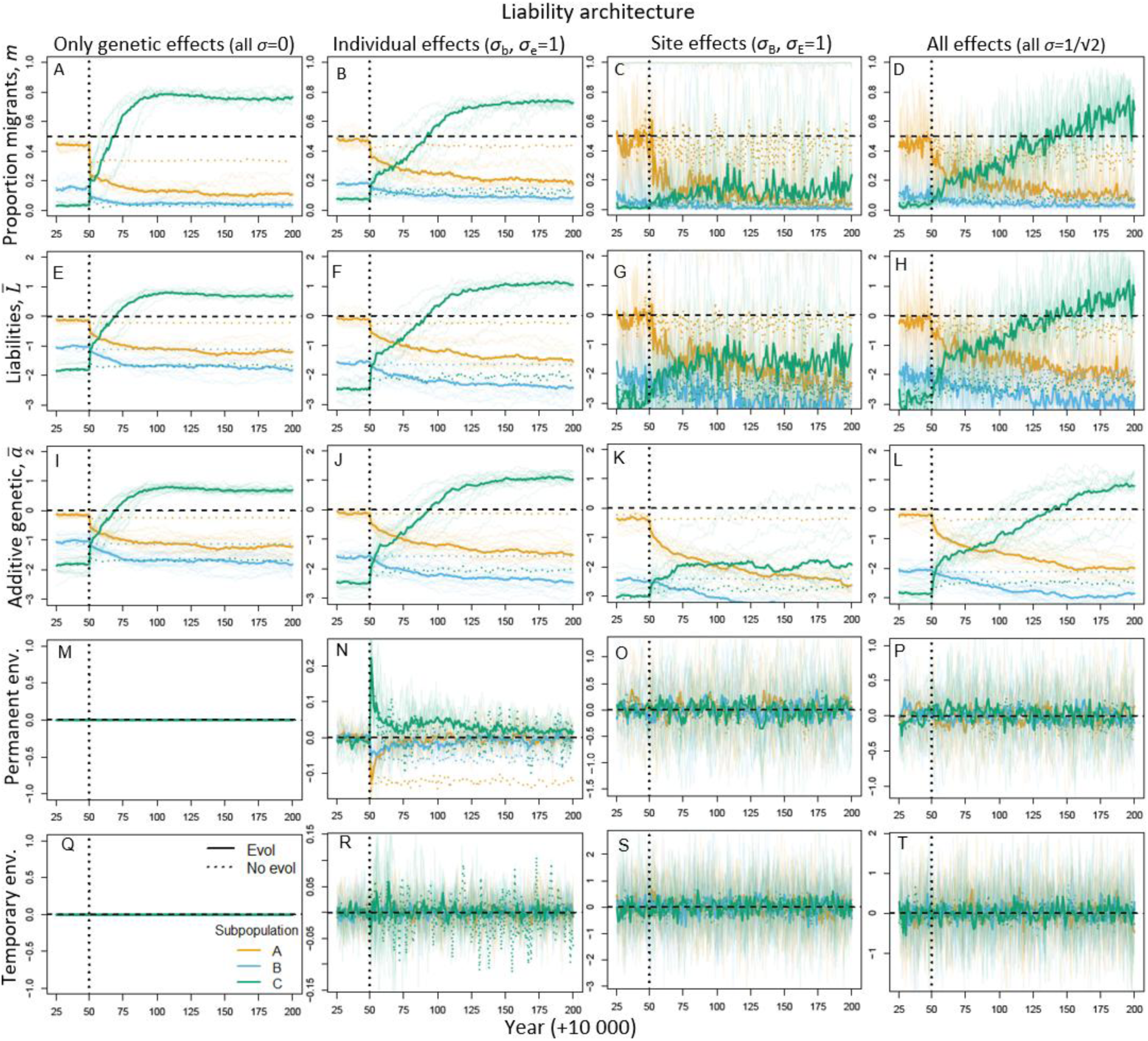
Evolutionary dynamics of seasonal migration following sudden environmental deterioration in site C from year 10050 (dotted vertical line) where the strength of non-breeding season density-dependence *γ*_C_=2, i.e. the end-point of the gradual deterioration scenario presented in main text. Columns show four liability architectures, comprising only genetic effects (column 1), permanent and temporary individual-specific environmental effects (column 2), permanent and temporary site-specific environmental effects (column 3), and all effects (column 4). Panels show trajectories for 150 years following onset of deterioration, for (A-D) proportion of migrants, (E-H) mean liabilities to migrate (*L̅*), (I-L) mean breeding values for liability (*a̅_j_*_,*t*_), (M-P) mean permanent environmental deviations (*σ*_B_*B̅*+*σ*_b_*b̅*), and (Q-T) mean temporary environmental deviations (*σ*_E_*E_j,t_*+*σ*_e_*e̅*). In all panels, orange, blue and green denote subpopulations A, B and C respectively. Thick lines show medians across 20 replicate simulations and opaque lines show individual realizations (20 in A-D, only 10 plotted in E-P for visual clarity). Horizontal dashed lines in A-H show the threshold value of zero, distinguishing liabilities that induce seasonal migration versus year-round residence, corresponding to 50/50 migration/residence (*m*=0.5) in I-L. Solid and dotted colored lines denote simulations with and without evolution respectively. In panels M and Q, all lines are on zero (with green printed last) because there are no environmental deviations given the liability architecture with only genetic effects. Note that y-axis ranges differ across columns in the two bottom rows.

Conversely, *b̅*_C,*t*_ first becomes strongly positive (even approaching 0.3), and then gradually decreases (Fig. S4N, green line). However, since it still takes several decades for subpopulation C to become sufficiently seasonally migratory, *b̅*_C,*t*_ remains positive and different from zero for a long time as micro-evolution of *a̅*_C,*t*_ progresses (Fig. S4J). As it becomes more migratory (through selection on both *a_i_* and *b_i_*), subpopulation C’s breeding population size grows, and consequently decreases the size of subpopulations A and B (Fig. S5A,B,D), as non-breeding season densities in sites A and B increase, strengthening density-dependent mortality (Fig. S5E-H). Thus, as for the main text scenario, evolution ‘rescues’ subpopulation C, but reduces population sizes for subpopulations A and B (Fig. S5I,J,L). As in the main text scenario, the architecture with only site-specific environmental deviations in most cases fails to evolve migration in subpopulation C (Fig. S4C), because slower evolution leads to protracted maladaptation decreasing population size enough that density-dependent selection for migration is relaxed. Here, subpopulation size in C increases during evolution (Fig. S5K) only because it allows subpopulations A and B to evolve full residence, lowering non-breeding season densities in C (Fig. S5G).

**Figure S5:**
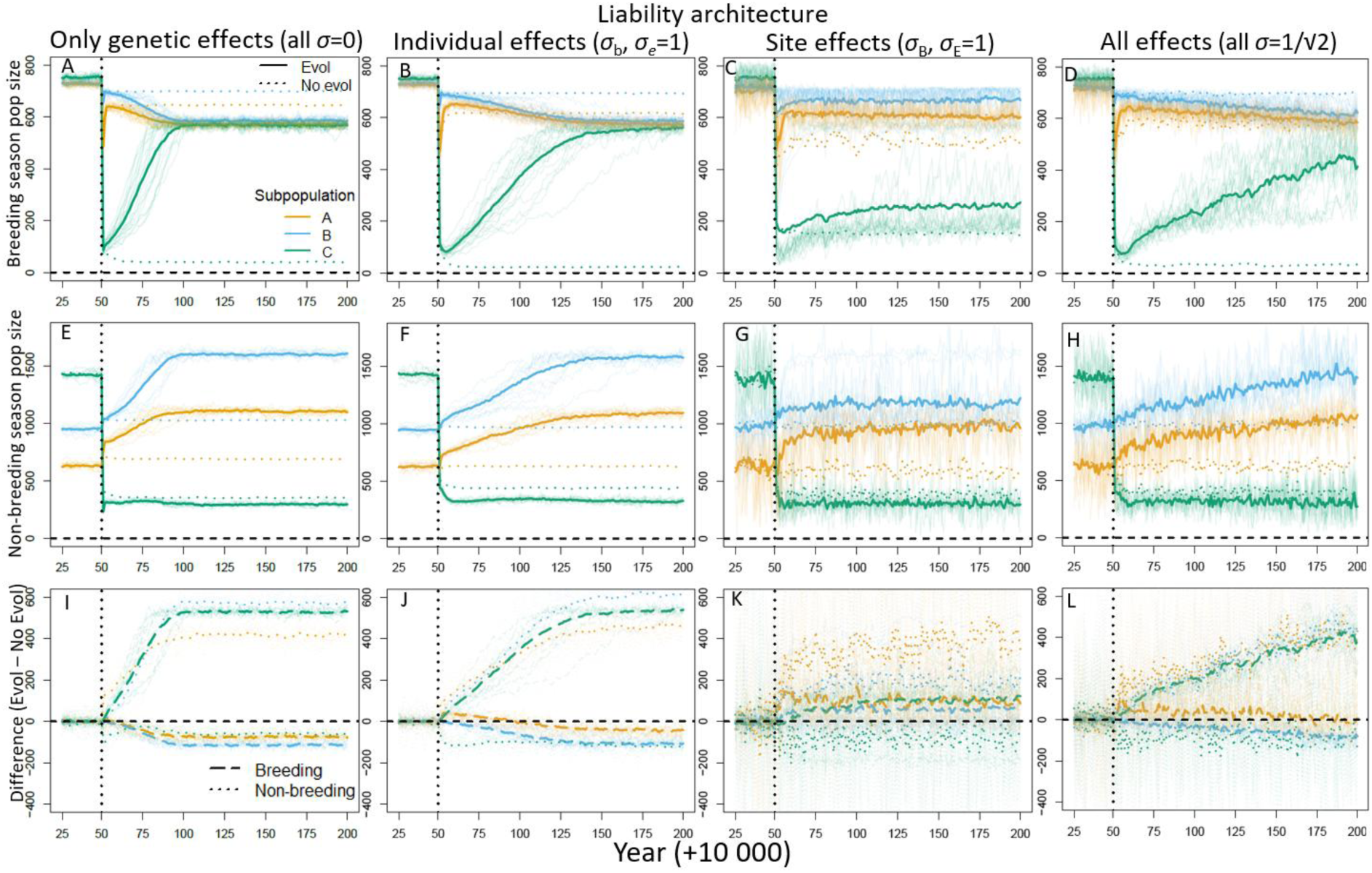
Spatio-seasonal population dynamics following sudden environmental deterioration in site C from year 10050 (dotted vertical line) where the strength of non-breeding season density-dependence *γ*_C_=2, i.e. the end-point of the gradual deterioration scenario presented in main text. Columns show four liability architectures, comprising only genetic effects (column 1), permanent and temporary individual-specific environmental effects (column 2), permanent and temporary site-specific effects (column 3), and all effects (column 4). Panels show trajectories over 150 years following onset of deterioration, for (A-D) breeding season adult population sizes, (E-H) non-breeding season population size for each site (i.e., number of individuals present, regardless of their subpopulation of origin), and (I-L) difference between local population sizes with vs without evolution (positive values indicate larger sizes with evolution than without). Orange, blue and green denote subpopulations A, B and C respectively. Thick lines show grand means across 20 replicate simulations and opaque lines show individual realizations (20 in panels A-D, only 10 plotted in E-L for visual clarity). In panels A-H, solid and dotted lines denote simulations respectively with and without evolution. In panels I-L, dashed and dotted lines denote differences for respectively breeding and non-breeding season population sizes.

#### ii) Gradual environmental deterioration, density-independent

We next enact gradual environmental deterioration by increasing non-breeding season density-*in*dependent mortality in site C. Specifically, we let the mortality rate of any individuals present deterministically increase by 0.01/year from 0.3 in year 10050 (which is typically less than the density-dependent mortality) to 0.5 in year 10070. The mortality probability for all individuals spending the non-breeding season in site C is then given by whichever is largest of the density-dependent mortality 1–*φ*_w,C,*t*_ and the density-independent mortality imposed by environmental deterioration. This parameterisation was chosen so that equilibrium metapopulation seasonal movements were broadly similar following environmental deterioration in both density-dependent and density-independent mortality (compare Fig. 2E-H with Fig. S6).

Here, evolutionary trajectories are broadly similar to the gradual density-dependent deterioration scenario presented in the main text for the liability architectures with only genetic effects and permanent and temporary individual-specific deviations (compare Fig. S7 with Fig. 3, first and second columns), although initial evolutionary responses in *a̅_j_*_,*t*_ tend to be slower (Fig. S7E,F vs. Fig. 3E,F). This is because the density-independent mortality rate in early years of deterioration in site C is typically less than the density-dependent mortality rate 1–*φ*_w,C,*t*_. The initial declines in breeding population sizes are therefore also less steep than under the main text scenario (Fig. S8 vs. Fig. 4). As deterioration progresses, the major difference from the density-dependent deterioration scenario presented in the main text occurs in the architecture with site-specific deviations. Given density-dependent deterioration, slower rates of evolution causing protracted maladaptation in subpopulation C decrease population sizes so much that density-dependent mortality is weakened, eventually reversing selection for migration and causing recanalization of residence (Fig. 3K, 4C). In contrast, density-independent deterioration causes sustained selection for migration in subpopulation C, such that a substantial proportion of replicate simulations (8/20) succeed in evolving migration (Fig. S7C), and breeding population sizes better recover (Fig. S8C).

**Figure S7:**
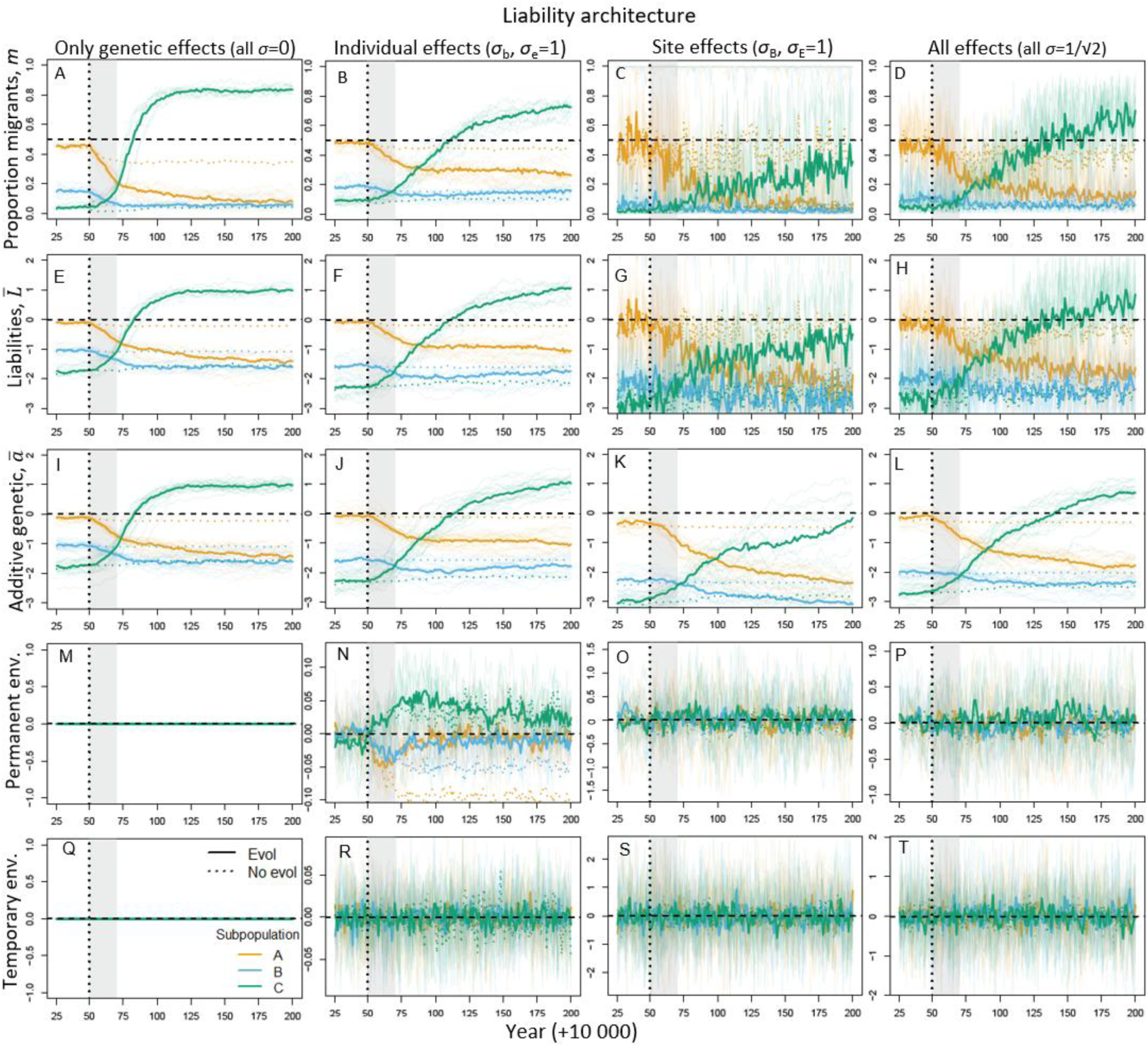
Evolutionary dynamics of seasonal migration during and following non-breeding season gradual density-independent environmental deterioration in site C starting from year 10050 (dotted vertical line) and lasting for 20 years (grey shaded area). Columns show four liability architectures, comprising only genetic effects (column 1), permanent and temporary individual-specific environmental effects (column 2), permanent and temporary site-specific environmental effects (column 3), and all effects (column 4). Panels show trajectories for 150 years following onset of deterioration, for (A-D) proportion of migrants, (E-H) mean liabilities to migrate (*L̅*), (I-L) mean breeding values for liability (*a̅*), (M-P) mean permanent environmental deviations (*σ*_B_*B̅*+*σ*_b_*b̅*), and (Q-T) mean temporary environmental deviations (*σ*_E_*E_j,t_*+*σ*e*e̅*). In all panels, orange, blue and green denote subpopulations A, B and C respectively. Thick lines show medians across 20 replicate simulations and opaque lines show individual realizations (20 in A-D, only 10 plotted in E-P for visual clarity). Horizontal dashed lines in A-H show the threshold value of zero, distinguishing liabilities that induce seasonal migration versus year-round residence, corresponding to 50/50 migration/residence (*m*=0.5) in I-L. Solid and dotted colored lines denote simulations with and without evolution respectively. In panels M and Q, all lines are on zero (with green printed last) because there are no environmental deviations given the liability architecture with only genetic effects. Note that y-axis ranges differ across columns in the two bottom rows.

**Figure S8:**
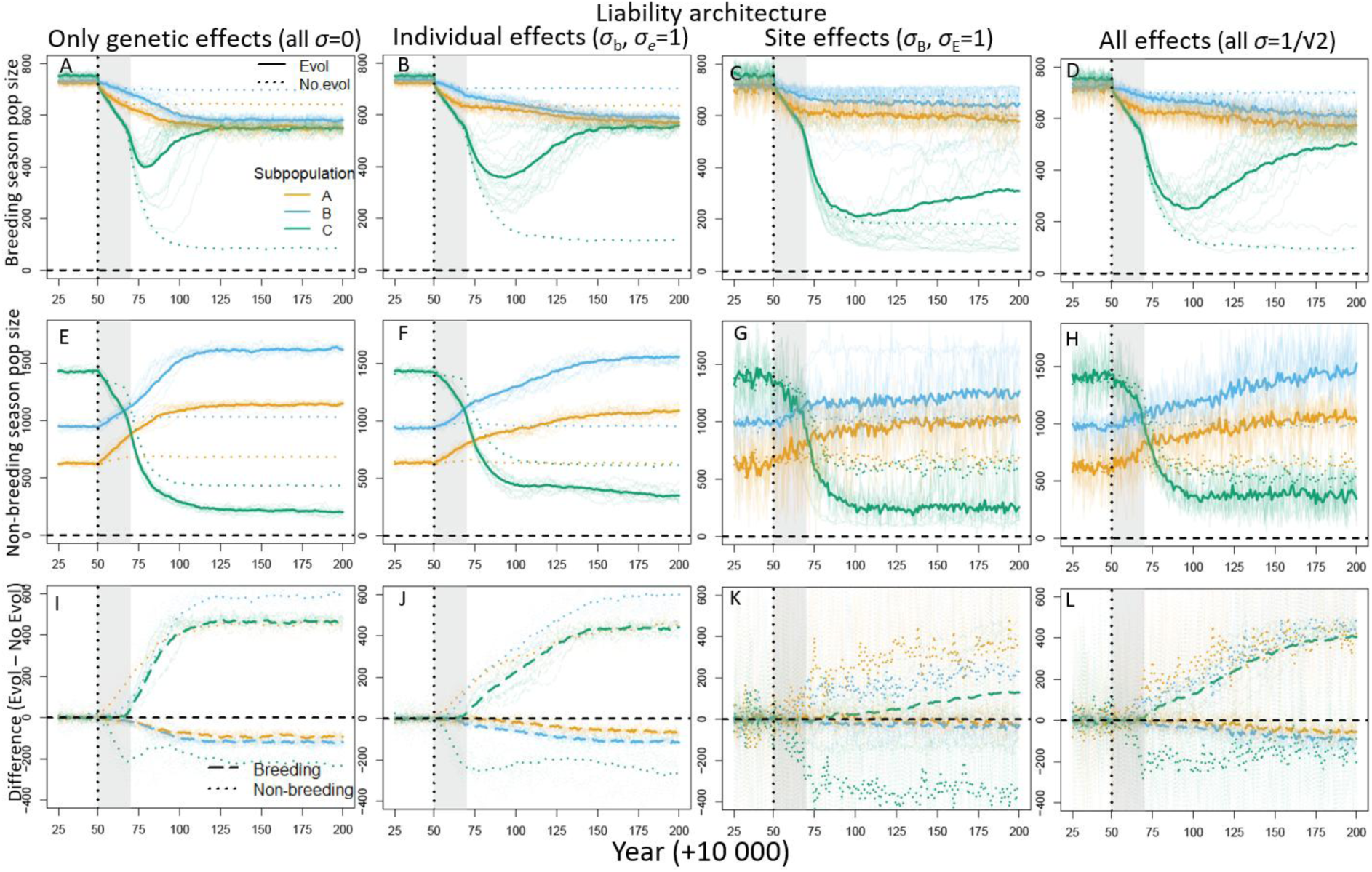
Spatio-seasonal population dynamics during and following non-breeding season gradual density-independent environmental deterioration in site C starting from year 10050 (dotted vertical line) and lasting for 20 years (grey shaded area). Columns show four liability architectures, comprising only genetic effects (column 1), permanent and temporary individual-specific environmental effects (column 2), permanent and temporary site-specific effects (column 3), and all effects (column 4). Panels show trajectories over 150 years following onset of deterioration, for (A-D) breeding season adult population sizes, (E-H) non-breeding season population size for each site (i.e., number of individuals present, regardless of their subpopulation of origin), and (I-L) difference between local population sizes with vs without evolution (positive values indicate larger sizes with evolution than without). Orange, blue and green denote subpopulations A, B and C respectively. Thick lines show grand means across 20 replicate simulations and opaque lines show individual realizations (20 in panels A-D, only 10 plotted in E-L for visual clarity). In panels A-H, solid and dotted lines denote simulations respectively with and without evolution. In panels I-L, dashed and dotted lines denote differences for respectively breeding and non-breeding season population sizes.

#### iii) Sudden environmental deterioration, density-independent

Finally, we consider the case of a sudden increase in density-independent mortality, which we implement by setting non-breeding season mortality probability in site C directly to 0.5 in year 10050 (i.e. the same as the end point for the gradual deterioration scenario).

This sudden environmental deterioration causes a less strong initial decrease in the size of subpopulation C than with a sudden increase in density-dependent mortality (compare Figs. S9-S10 with Figs. S3-S4). This is because of the high local population density at the time of the sudden environmental deterioration, which causes a very high density-dependent mortality rate. This rate thereafter declines as ensuing lower population sizes also lead to less density-dependent mortality, whereas the density-independent mortality rate remains the same even after the initial population reduction. Therefore, population recovery in subpopulation C is much slower when sudden environmental deterioration is enacted through density-independent than density-dependent mortality (Fig. S4A-D vs. Fig. S10A-D). A further consequence of environmental deterioration being enacted through density-independent rather than density-dependent mortality is that subpopulations A and B no longer experience a sudden population crash, then quickly recover, as in Fig. S4A-D. Rather, they experience a smaller initial crash, and then continue to gradually decline (Fig. S10A-D) as non-breeding season densities in sites A and B increase (Fig. S10E-H).

Despite these differences among environmental deterioration scenarios, our main conclusions regarding effects of different liability architectures still hold.

**Figure S9:**
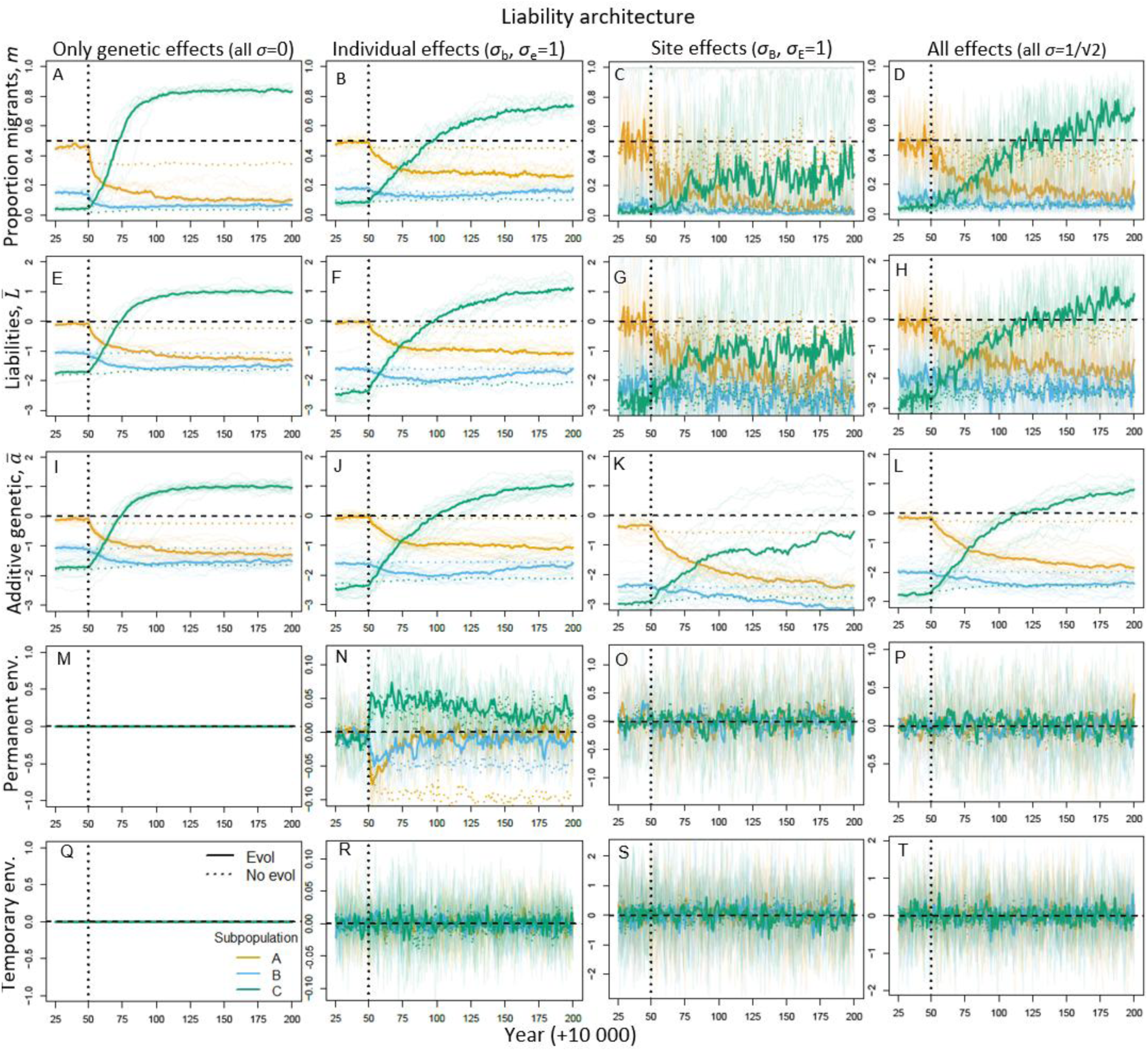
Evolutionary dynamics of seasonal migration following sudden density-independent environmental deterioration in site C from year 10050 (dotted vertical line) where non-breeding season mortality is set to 0.5, i.e. the end-point of the gradual deterioration scenario presented in Supplementary Material SM3ii. Columns show four liability architectures, comprising only genetic effects (column 1), permanent and temporary individual-specific environmental effects (column 2), permanent and temporary site-specific environmental effects (column 3), and all effects (column 4). Panels show trajectories for 150 years following deterioration, for (A-D) proportion of migrants, (E-H) mean liabilities to migrate (*L̅*), (I-L) mean breeding values for liability (*a̅*), (M-P) mean permanent environmental deviations (*σ*_B_*B̅*+*σ*_b_*b̅*), and (Q-T) mean temporary environmental deviations (*σ*_E_*E_j,t_*+*σ*_e_*e̅*). In all panels, orange, blue and green denote subpopulations A, B and C respectively. Thick lines show medians across 20 replicate simulations and opaque lines show individual realizations (20 in A-D, only 10 plotted in E-P for visual clarity). Horizontal dashed lines in A-H show the threshold value of zero, distinguishing liabilities that induce seasonal migration versus year-round residence, corresponding to 50/50 migration/residence (*m*=0.5) in I-L. Solid and dotted colored lines denote simulations with and without evolution respectively. In panels M and Q, all lines are on zero (with green printed last) because there are no environmental deviations given the liability architecture with only genetic effects. Note that y-axis ranges differ across columns in the two bottom rows.

**Figure S10:**
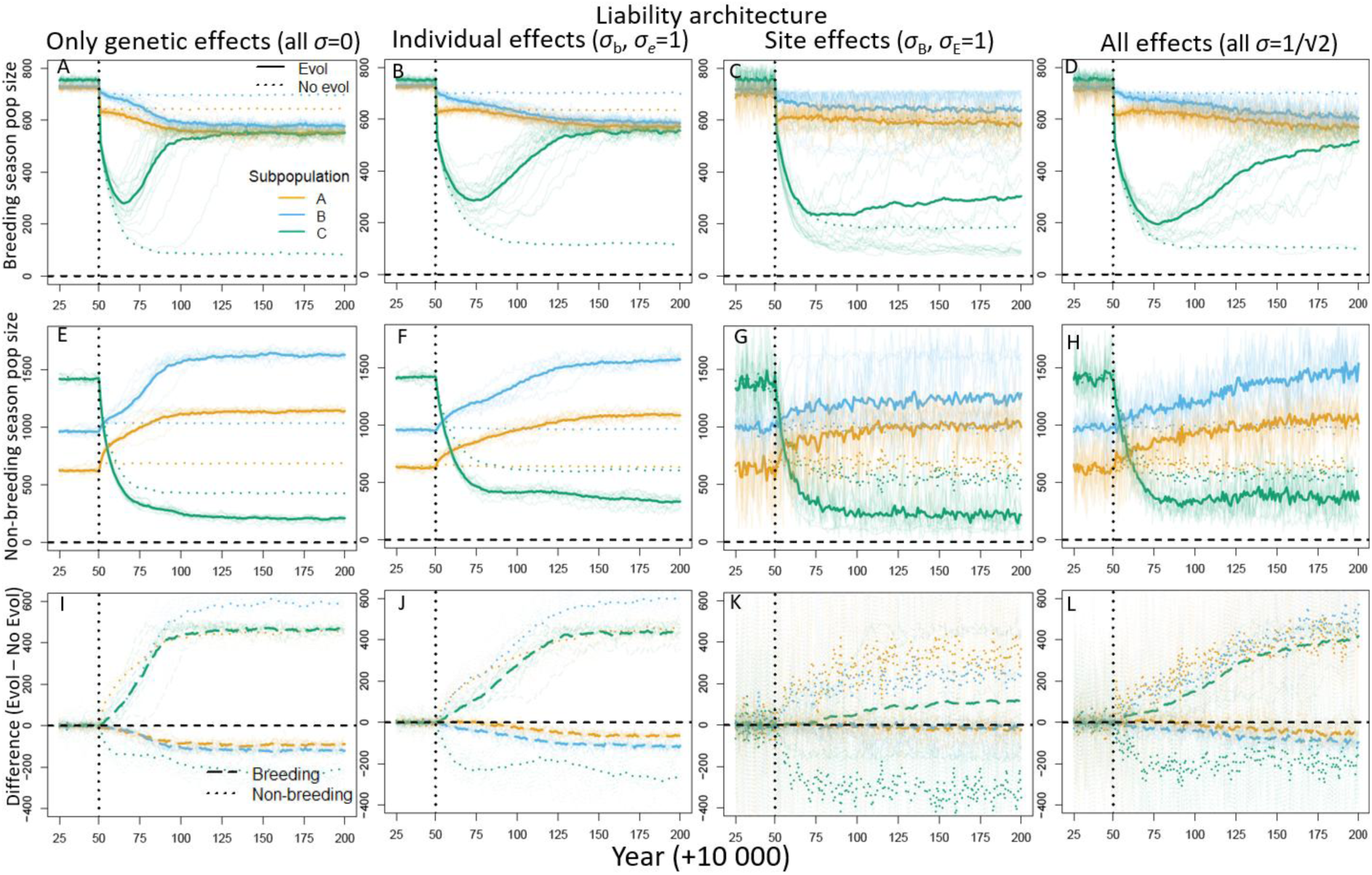
Spatio-seasonal population dynamics of seasonal migration following sudden density-independent environmental deterioration in site C from year 10050 (dotted vertical line) where non-breeding season mortality is set to 0.5, i.e. the end-point of the gradual deterioration scenario presented in Supplementary Material SM3ii. Columns show four liability architectures, comprising only genetic effects (column 1), permanent and temporary individual-specific environmental effects (column 2), permanent and temporary site-specific environmental effects (column 3), and all effects (column 4). Panels show trajectories for 150 years following deterioration, for (A-D) breeding season adult population sizes, (E-H) non-breeding season population size for each site (i.e., number of individuals present, regardless of their subpopulation of origin), and (I-L) difference between local population sizes with vs without evolution (positive values indicate larger sizes with evolution than without). Orange, blue and green denote subpopulations A, B and C respectively. Thick lines show grand means across 20 replicate simulations and opaque lines show individual realizations (20 in panels A-D, only 10 plotted in E-L for visual clarity). In panels A-H, solid and dotted lines denote simulations respectively with and without evolution. In panels I-L, dashed and dotted lines denote differences for respectively breeding and non-breeding season population sizes.

### SM 4: Transient emergence of phenotypic plasticity

In quantitative genetic threshold traits, the presence of any temporary environmental deviations in liability (here, *σ*_E_ or *σ*_e_>0) can cause phenotypic plasticity (i.e., expressing different alternative phenotypes in successive time steps), and this is intrinsically more likely when mean liabilities are closer to the threshold. During the course of adaptation to environmental deterioration, any evolution of mean breeding values towards the threshold then causes increased phenotypic plasticity, and subpopulations evolving away from the threshold causes decreased phenotypic plasticity (increased phenotypic canalization).

To quantify these changes in plasticity in seasonal migration versus residence over time in our simulations, we extracted individuals that lived ≥5 years and classified them as phenotypically plastic if they expressed both alternative phenotypes at least once, and fixed (canalized) otherwise. We then computed the frequencies of plastic, fixed resident and fixed migrant individuals alive within a given time window for each replicate (see Supplementary Material SM1). Here, choosing some minimum age is necessary to infer the phenotypic implications of population distributions of *L_i,j,k,t,t_*_0_, because individuals that only survive one or few years have no or little opportunity to express between-year plasticity. The cutoff age may bias observed proportions towards residence or migration due to cohort selection (e.g. Kendall et al. 2011), because maladapted individuals die earlier. However, decreasing (or increasing) the cutoff monotonically decreases (or increases) the frequencies of plastic individuals. Further, because of the relatively short expected lifespans emerging in our current simulations, results were similar if plasticity was instead measured as numbers of phenotypic switches made (e.g. Ugland et al. 2026).

As an illustrative example, we consider gradual density-dependent environmental deterioration (as in main text) and the liability architecture with permanent and temporary individual deviations (i.e., *σ*_b_ and *σ*_e_=1, Figs. 3-5, second column). Before onset of deterioration, subpopulation C contains predominantly (∼75%) canalized residents (green violins, Fig. S11A; see also Fig. 2B). Then, during adaptation to the new environmental regime, it passes through a more substantially partially migratory phase featuring increased phenotypic plasticity (Fig. S11B). Finally, it evolves towards canalized full migration (Fig. S11C; see also Fig. 2F), with an associated decrease in phenotypic plasticity (Fig. S11B). In contrast, in the initially highly plastic subpopulation A where *a*_A_ starts near the threshold (orange violins, Fig. S11B), evolution instead decreases phenotypic plasticity, in this case causing rapid canalization of residence (Fig. S11A). Similar patterns occur for other environmental deterioration scenarios and liability architectures (results not shown), except as mentioned for architectures without any temporary environmental deviations.

**Figure S11:**
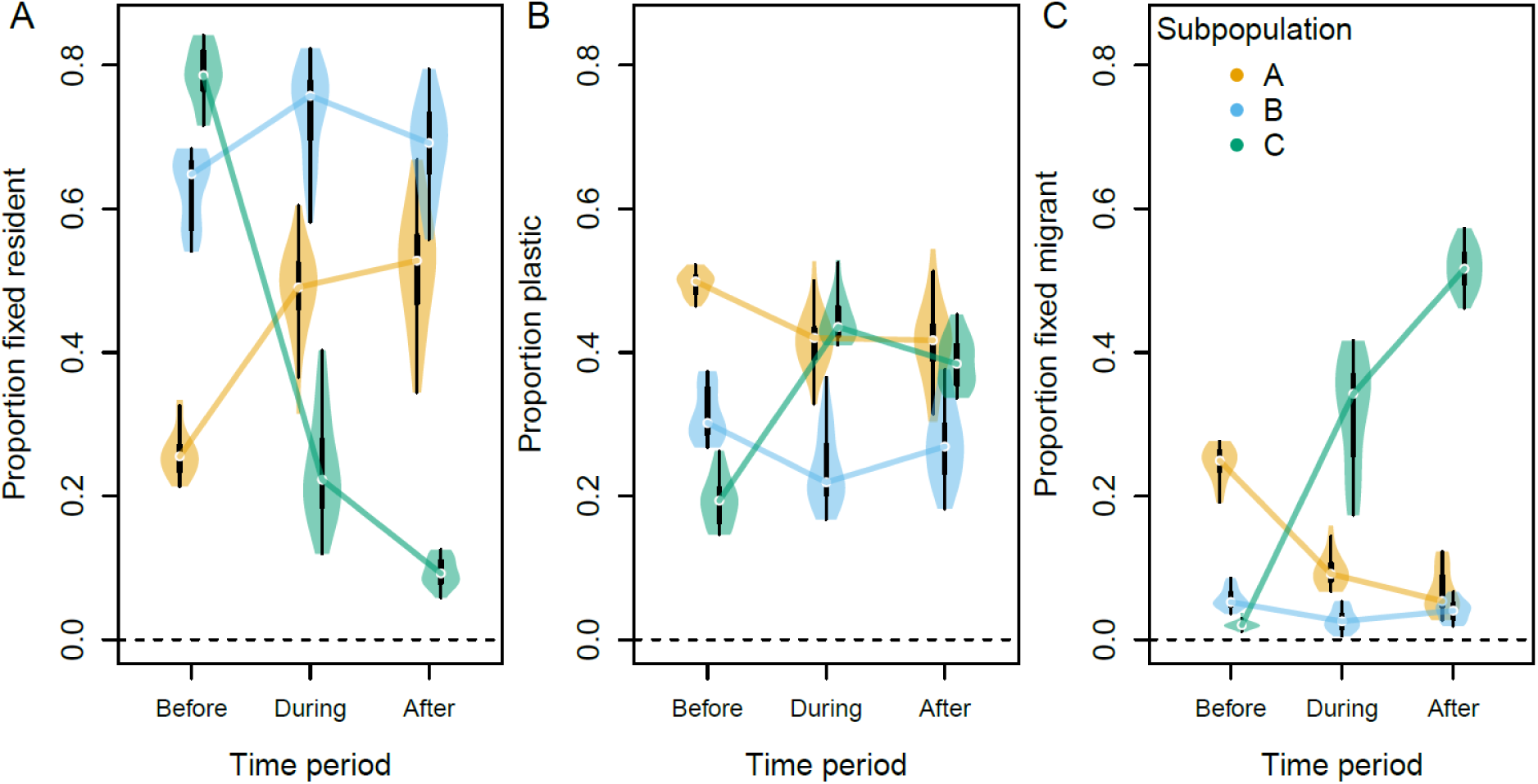
Proportions of (A) fixed resident, (B) phenotypically plastic, and (C) fixed migrant individuals in each subpopulation before, during and after adaptation to environmental deterioration in site C (x-axes). The liability architecture includes permanent and temporary individual-specific effects (*σ*_b_=1, *σ*_e_=1, as in second column of Figs. 3-4). Proportions are calculated across 20 years each (Before: 10001-10020; During: 10091-10110; After: 10181-10200), for individuals that lived for 5 years or more. Dots and thick bars in violins show median and interquartile range across 20 replicates. Orange, blue and green denote subpopulations A, B and C respectively.

Thus, despite no evolution of liability-scale reaction norms in our simulations (*σ*-parameters held constant throughout the duration of adaptation), the imposed environmental deterioration generates evolution of decreased phenotypic plasticity (increased canalization) in subpopulations A and B where liabilities evolve away from the threshold, and transient increases in phenotypic plasticity in subpopulation C that evolves towards and then beyond the threshold. This latter pattern echoes ‘plasticity-first’ evolution (c.f. Price et al. 2003; Schwander and Leimar 2011; Levis and Pfennig 2016; Kelly 2019), where environmental change initially increases phenotypic plasticity, then increases canalization, as evolution progresses. Such reductions in phenotypic plasticity during adaptation to a novel environment (also termed genetic assimilation, Scheiner and Levis 2021) has caused much debate (West-Eberhard 2003; Crispo 2007; Schwander and Leimar 2011). Although early work on genetic assimilation considered discrete phenotypes (Waddington 1953; Bateman 1959), most recent theory envisages traits that are continuously distributed on observed phenotypic scales (Lande 2009; Chevin and Lande 2010; Scheiner et al. 2020; Scheiner and Levis 2021; but see Raju et al. 2023). Here, assimilation typically occurs through genetic correlations between reaction norm elevations and slopes (Lande 2009), or through selection on slopes stemming from costs of plasticity (Chevin and Lande 2010; Scheiner et al. 2020). In our simulations, effective assimilation of a dichotomous trait occurs not due to any evolution of liability-scale reaction norms, or associated costs, nor because plasticity directly facilitates evolution. Indeed, the environmental effects reduce the heritability of liability to migrate, and consequently decrease the rate of evolution. Rather, the assimilation stems from intrinsic properties of dichotomous quantitative genetic traits (Reid and Acker 2022), contrasting with common claims that genetic assimilation requires costs of plasticity (Scheiner et al. 2017, 2020).

